# Gap junctions arbitrate binocular course control in flies

**DOI:** 10.1101/2023.05.31.543181

**Authors:** Victoria O. Pokusaeva, Roshan Satapathy, Olga Symonova, Maximilian Jösch

## Abstract

Animals utilize visual motion cues to maintain stability and navigate accurately. The optomotor response, a reflexive behavior for visual stabilization, has been used to study this visuomotor transformation. However, there is a disparity between the simplicity of this behavior and the intricate circuit components believed to govern it. Here we bridge this divide by exploring the course control repertoire in *Drosophila* and establishing a direct link between behavior and the underlying circuit motifs. Specifically, we demonstrate that visual motion information from both eyes plays a crucial role in movement control through bilateral interactions facilitated by gap junctions. These electrical interactions augment the classic stabilization behavior by inverting the response direction and the behavioral strategy. Our findings reveal how animals combine monocular motion cues to generate a variety of behaviors, determine the functional role of the circuit components, and show that gap junctions can mediate non-linear operations with a decisive role in animal behavior.

## Introduction

During visual course control, animals rely on optic flow as an important source of information about their self-motion and the structure of the environment. Optic flow patterns change stereotypically with each of the animal’s movements along its body axes. Thus, by extracting global motion patterns, an animal can generate a faithful inference of its self-motion and elicit stabilizing responses. The classic example of a visual course stabilizing response is the optomotor reflex^1^, a counteractive compensatory reaction observed in many animals^2^ that minimizes retinal slip during inadvertent movements. However, the accurate interpretation of visual motion cues is often equivocal. Distinct movements can result in similar local optic flow patterns, especially when the visual structure of the environment is nonuniform, as it usually is in natural environments. This challenges the ability of the visual system to instruct appropriate corrective motor commands since the same optic flow could be interpreted in multiple ways. Here, we use the fruit fly as a model to understand how such ambiguous motion cues are integrated binocularly across the visual field in order to instruct appropriate motor sequences.

In the fly’s brain, the circuit thought to be primarily involved in steering stabilizing responses is located in the posterior part of the lobula complex, called the lobula plate^3^. This circuit comprises a group of ∼30 direction-selective neurons in each hemisphere called lobula plate tangential cells (LPTCs)^3^. Different classes of LPTCs encode optic flow generated by self-motion about either the yaw, roll, or pitch axes^4^. One such class includes three yaw motion-sensitive horizontal system (HS) cells that respond to front-to-back (FtB) motion by changing their graded potential^5^ and two spiking neurons, H1 and H2 cells, that increase their firing rate to back-to-front motion^6^. Optogenetic and chemogenetic activation of HS-cells in *Drosophila* evoked directed head movement and flight turns^7–9^. Accordingly, silencing of HS neurons reduced head optomotor response, albeit without substantially affecting body turns^10^, suggesting that they are involved in gaze and course stabilization.

The sensitivity to a specific pattern of optic flow in the LPTCs arises primarily from the spatial organization of the direction-selective input onto these neurons^11^. Nevertheless, this specificity is further enhanced by the synaptic interactions between individual LPTCs within one or both hemispheres of the fly brain. Synaptic connections between LPTCs are mostly electrical^5,12–15^ and comprise ShakB gap junction channels^12,13,16,17^. These lateral electrical synapses mediate a direct flow of electrical current, allowing individual neurons to integrate motion information from visual areas outside their retinotopic dendritic inputs. Indeed, experimental and modeling studies suggest that gap junctions between ipsilateral LPTCs enable robust encoding of flow-field parameters by refining the structure of their spatial receptive fields^18^. Interestingly, these electrical connections have been shown to mediate binocular interactions too^13,14^. The behavioral role of this contralateral connectivity remains speculative, being suggested to be required to disambiguate from inadvertent translational and rotational movements in the horizontal plane. However, despite numerous predictions^13,17,19^, there is little direct evidence establishing the role of electrical connections in the steering responses generated by LPTCs.

In this study, we uncover the crucial role of binocular interactions in the fly’s course control system. Using a combination of behavioral, electrophysiological, and genetic approaches, we show that bilateral electrical connection plays a fundamental role in interpreting ego-motion-induced optic flow patterns, expanding our understanding of the computational roles of electrical synapses in sensorimotor transformations and behavioral control.

## Results

### Antagonistic behavioral responses to monocular optic-flow patterns

To explore the role of binocular integration in locomotion, we built a circular arena where flies walked freely, their position was tracked, and the body axis was used for closed-loop presentation of visual stimuli onto the arena’s roof (Figure 1A and see methods). When presenting wild-type CantonS flies with a radial sinusoidal grating (pinwheel), we reproduced the classic optomotor behavior, a strong and robust turning response in the direction of the stimulus (Figure 1B), composed of smooth turns and saccades (Figure 1C). Interestingly, we consistently observed saccades opposite to the direction of motion (anti-saccades) (Figures 1C,Di,E). These anti-saccades have been recently reported^20,21^, but their origin and functions remain unclear. Next, we decomposed the full-field rotational stimulus into back-to-front (BtF) and front-to-back (FtB) motion in either half of the visual field. In this modified pinwheel stimulus, only half of the pinwheel rotated while the other half remained static, centered on the fly’s body axis, allowing us to present either BtF or FtB motion to one of the two halves of the visual field. While BtF motion elicited a weak and transient turning response in the direction of motion (Figures 1Dii, 1E), FtB motion unexpectedly resulted in strong opposite turns (Figures 1Diii, 1E). Crucially, unlike a recent report of anti-directional turning behavior during optomotor response^21^, the anti-optomotor response to FtB motion that we observe (i) is initiated immediately after the motion onset, (ii) is sustained over the whole duration of the trial and (iii) shows little to no turning in the direction of rotation.

**Figure 1.**
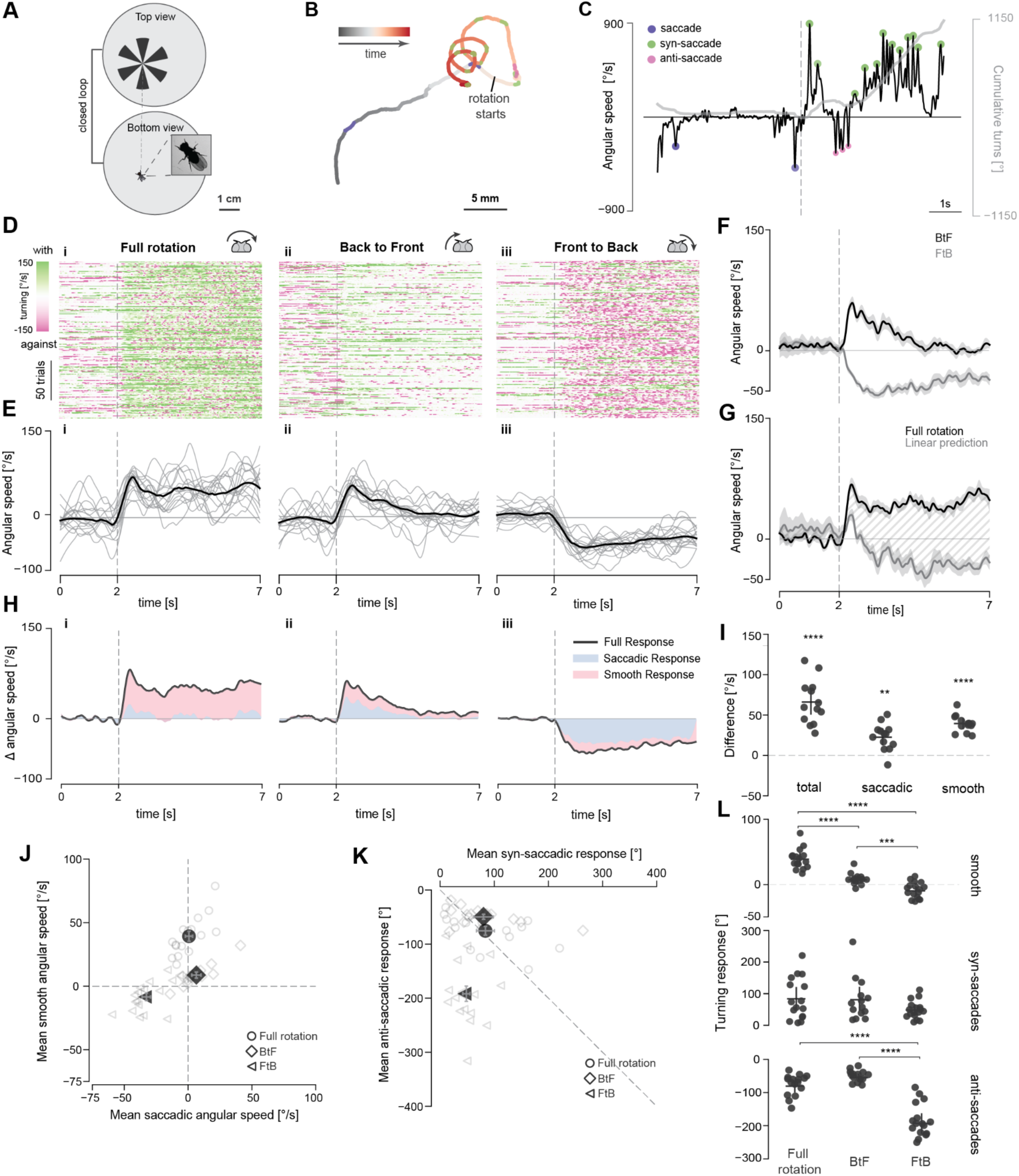
Differential binocular control of walking behavior in wild-type flies. (A) Schematic of the closed-loop behavioral setup. (B) Example trajectory of a CantonS fly during a typical trial. (C) Angular speed (black) and heading (gray) of the fly corresponding to the trial in (B) with syn- and anti-saccades highlighted. (D) Turning response of CantonS flies to (i) full-field, (ii) unilateral Back to Front (BtF) and (iii) unilateral Front to Back (FtB) rotation. Angular speed raster showing 200 randomly selected trials. Each row corresponds to one trial, each column to one frame (∼16 ms). (E) Mean angular speed per fly (gray) and across all flies (black). Dashed line indicates the beginning of the rotation. (F) Angular speed of CantonS flies in response to FtB and BtF rotation (mean ± SEM). (G) Comparison of the predicted and the actual mean angular speed in response to full-field rotation for CantonS flies. Shaded region shows the difference between predicted and actual response. (H) Stacked plot showing mean angular speed and the contribution of smooth and saccadic turning. (I) Mean prediction error across the trial period per fly. Bars: mean ± SEM. (J) Summary plot of the mean of smooth and saccadic angular speed across experiments. Empty and filled markers show the mean per fly and mean ± SEM, respectively. (K) Summary plot of the mean of syn-saccadic and anti-saccadic turns made. Empty and filled markers show the mean per fly and mean ± SEM, respectively. (L) Mean of smooth (top), syn-saccadic (middle) and anti-saccadic (bottom) turns made per trial. Bars: mean ± SEM. (I) One-sample t-test was applied to check if the error was significantly different from zero. (L) Two-sided Mann-Whitney U-test, ∗*p* < 0.05, ∗∗p < 0.01, ∗∗∗p < 0.001, ∗∗∗∗p < 0.0001. No asterisk: not significant. The number of flies and exact *p*-values for each experiment are listed in Suppl. Table1.

Since the full-field rotation of the pinwheel can be decomposed into FtB and BtF motion, we expected that the fly’s response to the full-field rotation would be a simple summation of its response to FtB and BtF motion. However, the observed full-field response is significantly higher than our linear prediction (Figures 1F, 1G), suggesting that the course control system of the fly integrates visual information from the two halves of the visual field in a non-linear manner. To determine the source of this non-linear behavioral response, we characterized the difference in behavioral properties across the three stimuli by separating the smooth and saccadic turning responses for each trial (see methods). While the classic optomotor response to full-field rotation is primarily composed of smooth turns (Figure 1Hi), the turning response to BtF motion has an equal contribution of both smooth and saccadic turning (Figure 1Hii). Strikingly, the anti-optomotor turning in response to FtB motion is predominantly saccadic (Figure 1Hiii). Accordingly, in response to FtB motion, the locomotion path is straighter (Figure S1A). Thus, flies alter the direction as well as the nature of their turning response for different types of motion, which accounts for the significant prediction errors for both smooth and saccadic turning (Figure 1I). The relative contribution of smooth and saccadic turning is evident across flies for each stimulus (Figure 1J,1L). Interestingly, although the number of anti-saccades per trial is higher (Figure S1B), syn- and anti-saccades contributed equally to the full-field optomotor responses (Figure 1K) due to larger syn-saccadic turns (Figures S1C, S1D). In summary, wild-type flies show opposite responses to unilateral FtB and BtF motion, indicating that they interpret these motions differently. Furthermore, these responses are integrated non-linearly to generate the classic optomotor response to full-field rotation.

### LPTCs are required for binocular control of walking

To define the circuitry instructing non-linear binocular behaviors, we focused on the lobula plate tangential cells (LPTC), a network of neurons sensitive to wide-field motion and thought to influence optomotor response in flies^13,22^. We targeted all and only horizontal system (HS) and vertical system (VS) neurons, using the VT058487-GAL4 line crossed to tsh-GAL80, a GAL4 repressor expressed in the ventral nerve cord, and silenced them by expression of the inward rectifying potassium channel Kir2.1 (Figure 2A). Surprisingly, silencing all HS and VS neurons did not abolish the classic optomotor response (Figure 2C), in spite of a decrease in the overall strength. However, the response to FtB motion was markedly different, with flies showing an early transient response against the direction of motion, followed by a late sustained response in the direction of motion (Figure 2B). More importantly, in flies with HS and VS neurons silenced, the response to full-field rotation matches the linear prediction (Figure 2C), as demonstrated by the significantly decreased prediction error compared to UAS-Kir2.1 control flies (Figure 2F-G).

**Figure 2.**
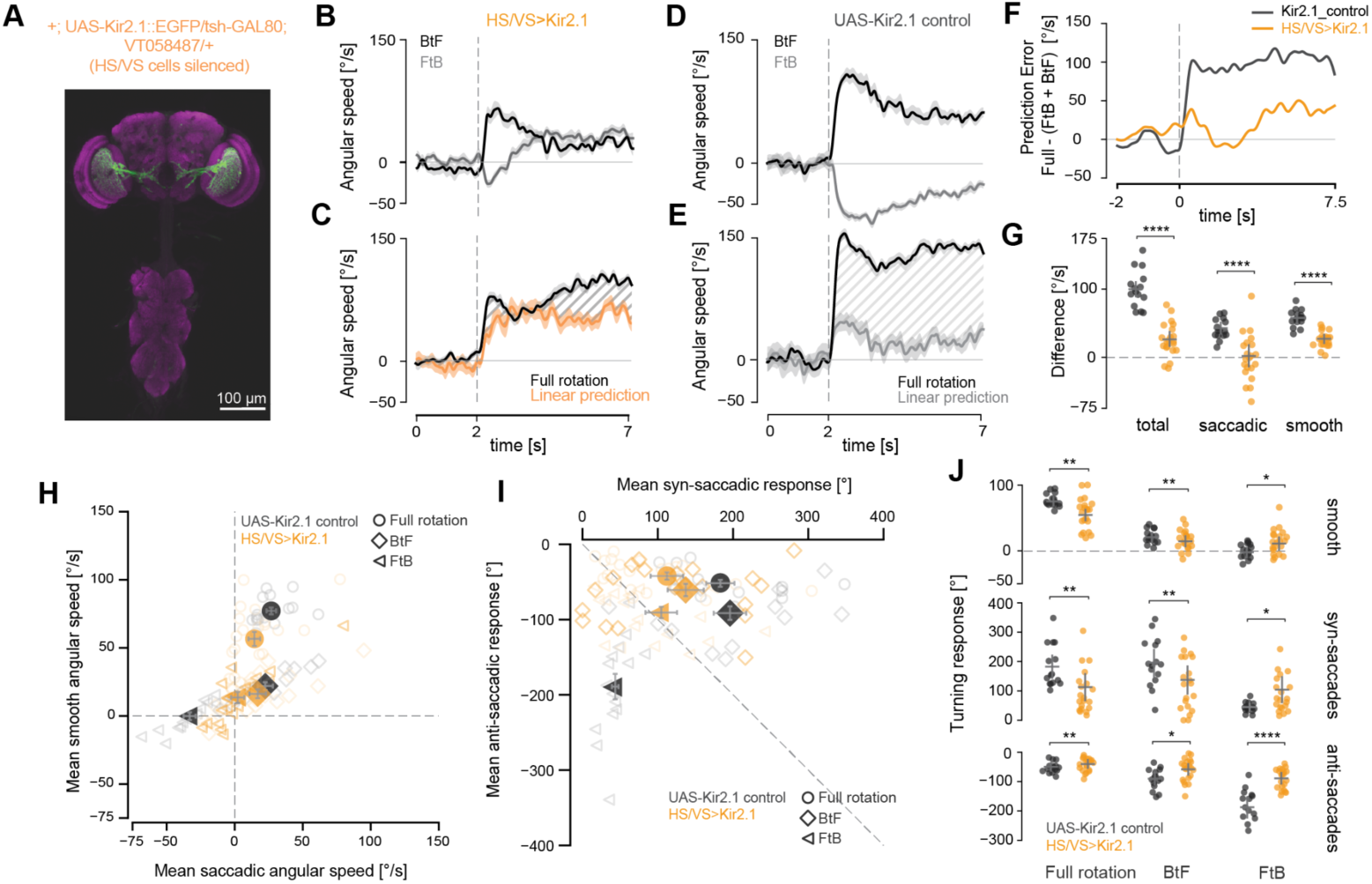
LPTCs are required for binocular control of walking. (A) Maximum z-projection of Kir2.1::EGFP expression by the VT058487-GAL4 line, used to silence HS and VS cells. The flies also carried a tsh-GAL80 transgene to eliminate GAL4 activity in the ventral nerve cord. (B) Angular speed of HS & VS-silenced flies in response to FtB and BtF rotation (mean±SEM). (C) Comparison of the predicted and the actual mean angular speed in response to full-field rotation for HS & VS-silenced flies. Shaded region shows the difference between predicted and actual response. (D) Same as in (B) but for UAS-Kir2.1 control flies. (E) Same as in (C) but for UAS-Kir2.1 control flies. (F) Time series of mean prediction error. (G) Mean prediction error across the trial period per fly. Bars: mean± SEM. (H) Mean of smooth and saccadic angular speed across a trial. Empty and filled markers show the mean per fly and mean ± SEM for each genotype, respectively. (I) Mean of syn-saccadic and anti-saccadic turns made in one trial. Empty and filled markers show the mean per fly and mean ± SEM for each genotype, respectively. (J) Mean of smooth (top), syn-saccadic (middle) and anti-saccadic (bottom) turns made per trial. Bars: mean± SEM. (G and J) Two-sided Mann-Whitney U test, ∗*p* < 0.05, ∗∗p < 0.01, ∗∗∗p < 0.001, ∗∗∗∗p < 0.0001. No asterisk: not significant. The number of flies per genotype and exact *p*-values for each experiment are listed in Suppl. Table1.

Changes in the turning dynamics of FtB response cause the linearization of optomotor response in flies with silenced HS and VS neurons. While the FtB response in control or wild-type flies is dominated by anti-saccades (Figures 1H,1K, 2H,2I, S2F), silencing of HS and VS neurons results in a decrease in anti-saccadic response and an increase in syn-saccadic response (Figures 2H-J, S2B). This causes the overall response to become smooth (Figure S2F), further demonstrated by a decrease in path straightness (Figure S2A). Consequently, the prediction error for saccadic turning is nearly zero in flies with silenced HS and VS neurons (Figure 2G).

While our experiments imply that HS neurons are responsible for generating anti-saccades in response to FtB motion, this conclusion is challenged by previous findings. HS cells are known to be excited by FtB motion^5^, and their unilateral activation has been shown to drive behavioral responses in the direction of movement^7,8^. Yet, wild-type flies turn against the direction of motion in response to FtB motion, strongly suggesting the involvement of other wide-field direction selective inputs. H2, a class of spiking LPTC that responds to horizontal BtF motion, is a possible candidate. Previous work in blowflies has shown that these neurons are electrically connected to HSE cells in the contralateral hemisphere and are their primary source of contralateral input^13^. Biophysically, this binocular interaction was shown to be nonlinear and mediated by electrical synapses^13^, suggesting that this specific electrical connectivity could mediate the putative non-linear binocular interactions we inferred from the behavioral experiments. We set out to test this hypothesis by genetically disrupting electrical synapses between LPTCs.

### Validation of a new inducible *shakB* mutant

The role of electrical synapses in *Drosophila* neural circuits is a topic of extensive research. However, genetic manipulation of gap junctions, the molecular substrates of electrical synapses, remains challenging. Widely used EMS-induced mutant lines, such as *shakB*^2^, offer the advantage of robust gene inactivation but frequently carry background mutations. Moreover, attempts at cell-specific inactivation of gap junctions in the fly visual system have been met with limited success^16^. Consequently, pinpointing the contribution of electrical synapses within visual neural circuits has remained elusive.

To attempt a detailed description of the role of electrical synapses in LPTCs while avoiding the pitfalls mentioned above, we generated transgenic animals that carry the FlpStop-cassette^23^ inside the innexin known to mediate LPTC electrical connectivity, *shakB*^16^ (Figures 3A,B, S3A). This approach enables us to generate both full and cell-specific mutants while maintaining an isogenic background, greatly facilitating the interpretation of behavioral experiments. FlpStopNDshakB (FlpND, non-disruptive orientation) and FlpStopDshakB (FlpD, disruptive orientation) flies were created by integrating FlpStop-cassette into an intronic MiMIC insertion between exons 5 and 6. This insertion allows the inactivation of 6 out of 8 isoforms of the ShakB protein, leaving isoforms *shakB-RA* and *shakB-RE* intact (Figure 3B). Notably, the widely-used *shakB*^2^ mutant carries a null mutation in 5 isoforms, leaving *shakB-RF* undisrupted in addition to *shakB-RA* and *shakB-RE*.

**Figure 3.**
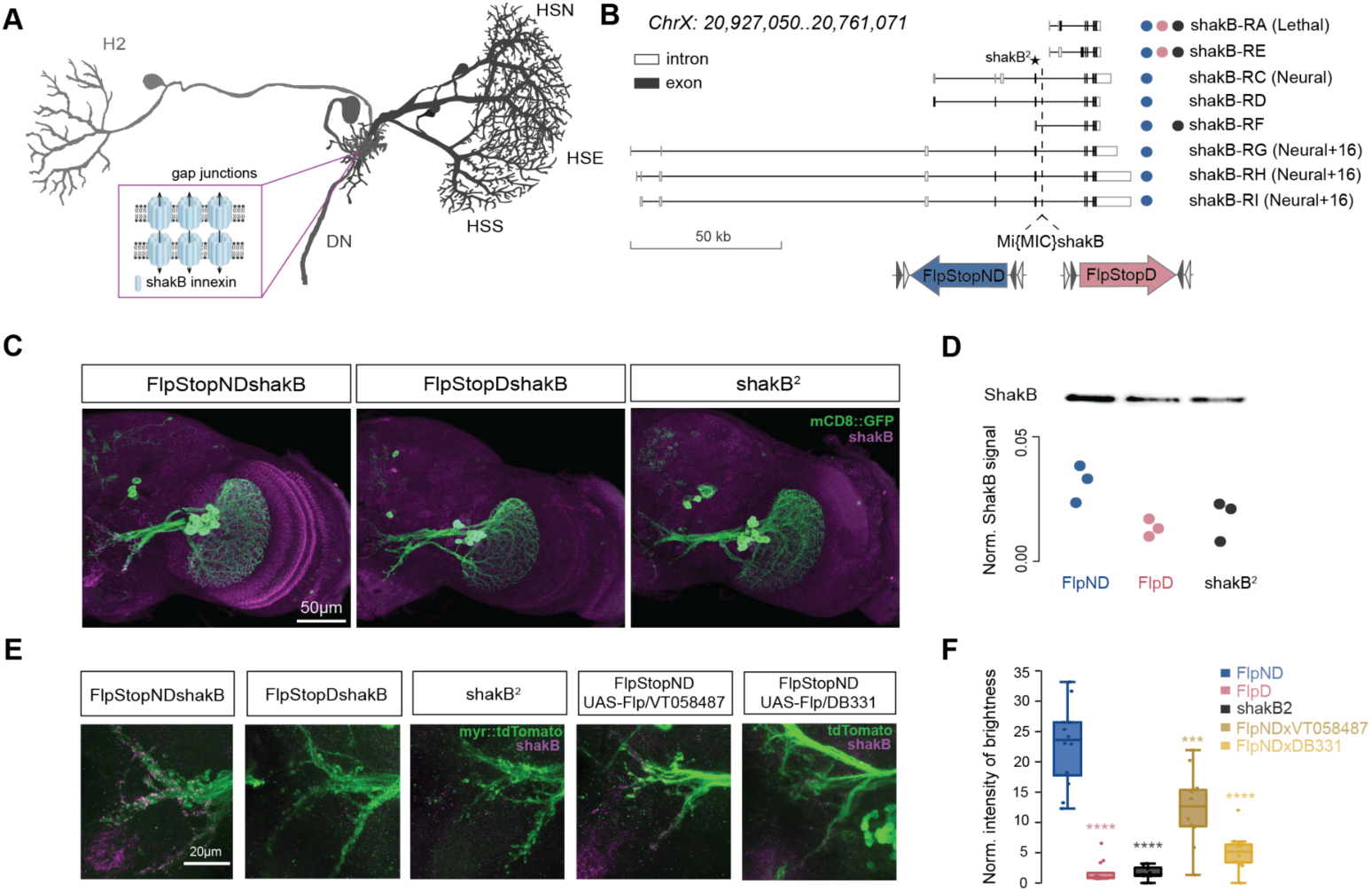
FlpStop *shakB* interventions reveal ShakB protein stability. (A) Schematic of known axonal gap junctions in horizontal system LPTCs. Pairs of ShakB innexins form transmembrane channels that enable bidirectional flow of electrical current between two adjacent neurons. (B) Gene map of *shakB* isoforms. Nonsense mutation carried by *shakB*^2^ flies is indicated with a star. Dashed line indicates the position of the intronic MiMIC cassette used to integrate the FlpStop cassette in disruptive (FlpD, magenta) and non-disruptive (FlpND, blue) orientations. The circles indicate isoforms that are intact in FlpND (blue), FlpD (magenta) and *shakB*^2^ (black) flies. (C) Immunostainings for the gap junction protein ShakB in the optic lobe of wild-type line FlpND and two mutant lines FlpD and *shakB*^2^. (D) The western blot analysis of protein extracts from fly brain tissue using antibodies against ShakB. ShakB protein signal was normalized to the total amount of the protein sample per line. (E) LPTCs axonal staining of ShakB protein in wild-type (FlpND), full mutant (FlpD and shakB^2^) and induced mutant (VT058487-Gal4 and DB331-Gal4) flies. (F) Quantification of the ShakB signal in LPTC axons (n = 6 brains per genotype, each hemisphere was quantified independently). All the mutant genotypes were compared to wild-type (FlpND) using two-sided Mann-Whitney U test, ∗∗∗p < 0.001, ∗∗∗∗p < 0.0001. The number of recorded cells and exact *p*-values for each experiment are listed in Suppl. Table1.

We observed a significant reduction in the total amount of ShakB protein in the brain of FlpD flies compared to FlpND flies (Figure 3C, 3D). Recent studies suggest that cell-specific inactivation of ShakB protein using driver lines selective to LPTCs is ineffective^16^. We hypothesized that it might result from early expression onset and a slow turnover rate of innexins in these neurons. The potential influence of developmental timing on the efficiency of cell-specific inactivation of *shakB* prompted us to trigger the inversion of FlpStop-cassette using two LPTC-specific driver lines - DB331-Gal4 and VT058487-Gal4. DB331-Gal4 line initiated Flp-mediated cassette inversion in LPTCs at around pupal stage P9 while VT058487-Gal4 at around P12 (Figure S3C). ShakB immunolabelling in LPTC axons showed that DB331-Gal4 induced a stronger knock-down of *shakB* than VT058487-Gal4 driver line (Figure 3E, 3F), indicating that the timing of gene inactivation is a critical determinant of the phenotype.

Loss of gap junction may severely affect the development of brain tissue^24^. Therefore, it is crucial to rule out developmental phenotypes to avoid misinterpreting physiological and behavioral experiments in the newly generated mutant flies. To account for the potential effects of *shakB* disruption on the expression of other proteins in the fly brain, we performed the proteomic analysis of brain tissue in FlpND, FlpD, and *shakB*^2^ flies. Out of 5755 proteins identified, only 7 were upregulated in FlpD flies, while 15 were up- and 26 were downregulated in *shakB*^2^ flies (Figure S4A, S4B). The higher number of affected proteins in *shakB*^2^ flies is probably due to its distinct genetic background to FlpND control flies but could also hint at additional extraneous mutations. Proteins Acbp2, Primo-1, and CG31345 were significantly up-regulated in both FlpD and *shakB*^2^ flies, indicating their functional connection with ShakB protein.

Gap junctions were previously shown to be involved in refining neuronal morphology^24^ and controlling the formation of chemical synapses^25^. To identify potential differences in the morphology of HS cells in wild-type and mutant flies, we fluorescently labeled individual HS cells using the SPARC technique^26^. The confocal image stacks were used for 3D reconstructions of neurons and then for morphology analysis. We observed no significant differences in dendritic branching and volume between HS cells in wild-type and mutant flies (Figure S5). To label postsynaptic partners of HS cells in mutant and wild-type flies, we used the trans-Tango technique^27^. The postsynaptic partners of HS cells were detected in the optic lobe, in the posterior slope, and in the ventral nerve cord (Figure S6). All the postsynaptic partners of HS cells observed in the wild-type flies were also observed in FlpD flies (Figure S6). Meanwhile, the trans-synaptic tracing in *shakB*^2^ flies revealed fewer synaptic partners of HS cells, with higher labeling variability suggesting that chemical synaptic connectivity might be altered in *shakB*^2^ mutants.

Overall, we show that the newly developed FlpStop-based system allows efficient inactivation of ShakB protein with little-to-no influence on the proteome as well as the morphology and formation of synaptic connections of HS cells. Furthermore, cell-specific inactivation of ShakB protein is driver-line-dependent and can be efficiently achieved only by the inversion of FlpStop cassette at early pupal stages.

### Loss of gap junctions does not abolish direction-selective responses

To characterize the passive membrane properties and visual responses of HS cells lacking gap junctions, we used *in vivo* whole-cell patch clamp recordings in tethered flies (Figure 4A)^15^. We first confirmed that integrating the FlpStop-cassette in non-disruptive orientation does not affect the direction-selective responses of HS cells and can be used as a wild-type control (Figures 4C, S7A, S7B).

**Figure 4.**
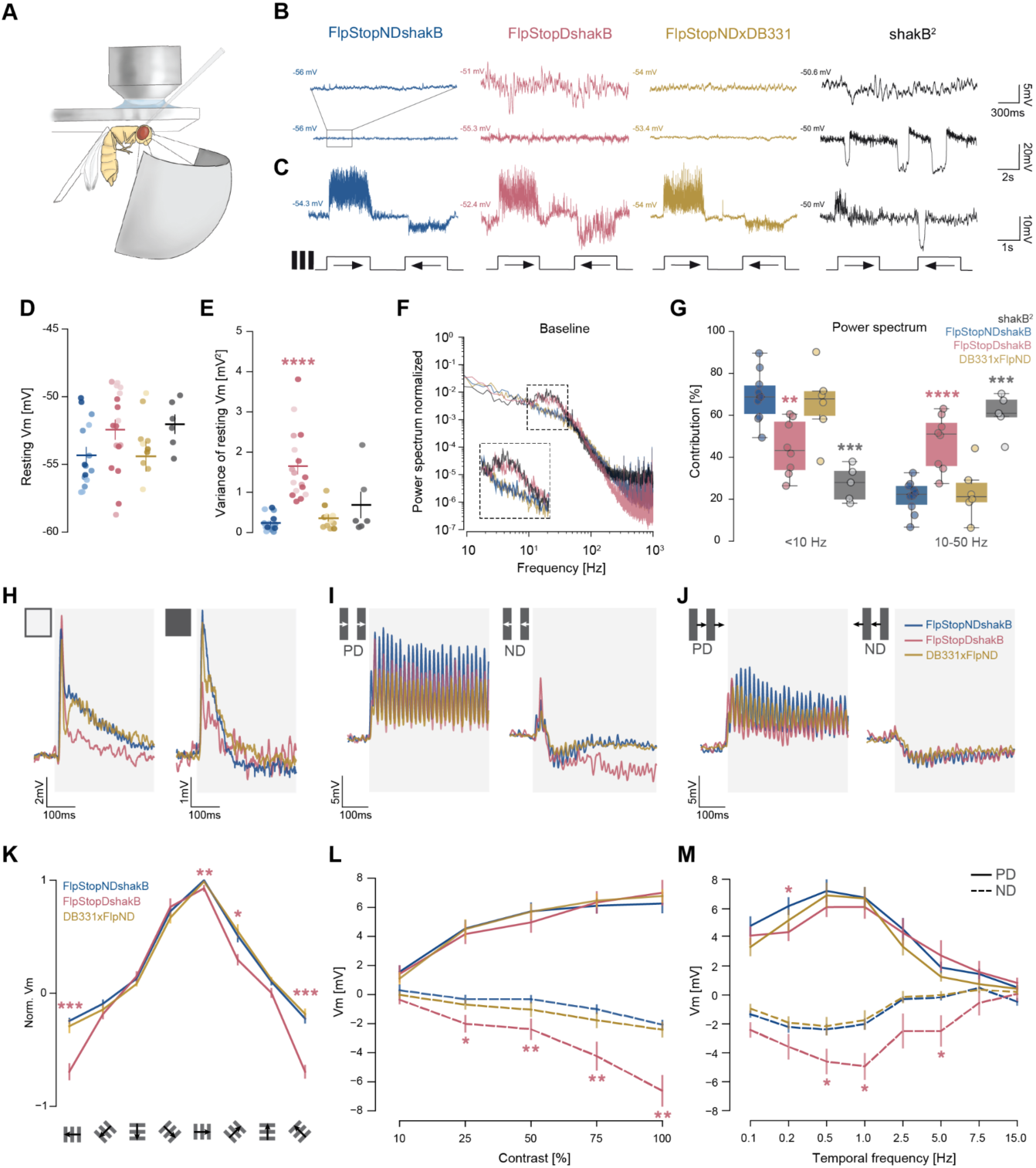
Visual responses and passive membrane properties of HS cells lacking gap junctions. (A) Set-up for *in vivo* whole-cell patch clamp recordings in tethered flies. (B) Example traces of membrane potential of HS cells while at rest and (C) during direction-selective responses. (D) Resting membrane potential of individual HS cells (mean± SEM). The saturation of the color depicts distinct HS types: the highest - HSN, the middle - HSE, the lowest - HSS (cell types were not identified for *shakB*^2^). (E) Variance of resting membrane potential (mean± SEM) as in (D). (F) Normalized power spectrum of baseline membrane potential in HS cells. (G) Contributions of low-range (<10 Hz) and mid-range (10-50 Hz) frequencies to the total power. (H) Average response traces of HS cells during the first 0.5 s after the onset of the ON/OFF flash. (I) Average response traces of HS cells to drifting ON-edges moving in PD and ND. (J) Same as (H), but for OFF-edges. (K) Normalized average voltage changes during 2 s presentation of square-wave gratings moving in 8 different directions (mean± SEM). (L) Average responses of HS cells during 2 s presentation of gratings with different contrast moving in PD and ND (1 Hz temporal frequency, mean± SEM). (M) Average responses of HS cells during 2 s of presenting gratings moving with different temporal frequencies (mean± SEM). (D), (E), (G), (K-M), all the mutant genotypes were compared to wild-type (FlpND) using two-sided Mann-Whitney U test, ∗*p* < 0.05, ∗∗p < 0.01, ∗∗∗p < 0.001. No asterisk: not significant. The number of recorded cells and exact *p*-values for each experiment are listed in Suppl. Table1.

On average, the resting membrane potential of HS cells in the FlpD mutant was higher and more variable than FlpND, consistent with observations made for the *shakB*^2^ mutant (Figures 4B, 4D, 4E). The membrane potential of HS cells in FlpD flies displayed spontaneous fluctuations (Figures 4B, 4F-G), similar to those described in *shakB*^2^ mutant flies^16^. Interestingly, while we detected fast β-oscillations of a frequency band similar to *shakB*^2^, we did not observe strongly hyperpolarizing ultraslow waves in FlpD flies (Figure 4B, 4G). Importantly, we observed membrane fluctuations in flies with cell-specific inactivation of *shakB* in LPTCs only in the rare events where dye coupling was fully abolished (Figure 4B, S7C), suggesting that only complete inactivation, and not a reduction of gap junction protein, can induce the membrane oscillations.

When presented with full-field flashes, ON-transient responses of HS cells in all animals had similar amplitudes, while OFF-transient responses were reduced in FlpD flies (Figure 4H, S7D,E). However, the reduction in OFF-transients was significantly smaller than what was previously reported for *shakB*^2^ flies^16^. In addition, we observed strong direction-selective responses in both mutant and wild-type flies in response to bright ON and dark OFF edges traveling at a velocity of 14°/s (Figures 4I-J, S7F-G). These results suggest that the loss of gap junctions in FlpD flies partially affects the visual processing of light decrement signals.

Moreover, HS cells in FlpD mutant flies did not show the reduced amplitude of direction-selective responses previously described for *shakB*^2^ flies^16^. On the contrary, we observed enhanced hyperpolarizing responses to gratings moving in null-direction (ND) in FlpD mutant flies but not in cell-specific DB331xFlpNDshakB flies. Interestingly, while ON-edge moving in ND elicited stronger hyperpolarizing responses in FlpD flies, the amplitude of responses to OFF-edge was similar across genotypes (Figures 4I-4J). It suggests that enhanced hyperpolarization is not a cell-intrinsic phenomenon. Finally, direction (Figure 4K), contrast (Figures 4L, S7H), and frequency tuning (Figures 4M, S7I) show that enhanced hyperpolarization do not affect the overall tuning properties of HS cells in FlpD flies.

Altogether, the analysis of electrophysiological properties shows that the membrane potential of HS cells in FlpD flies exhibits fluctuations that do not affect the tuning of direction-selective responses in these cells. Nevertheless, the loss of gap junctions increases hyperpolarization amplitude in response to ND motion.

### Electrical synapses shape HS cell receptive fields

To investigate the impact of gap junctions on the response properties mediated by the LPTC network, we conducted a thorough analysis of the receptive fields (RF) of HS cells in both FlpND and FlpD flies. As done previously^5^, we used a local moving spot that scanned the visual field in four cardinal directions to determine their local direction-selective responses across the fly’s bilateral visual field (140° in azimuth and 80° in elevation). The screen’s size and its dorsally displaced positioning did not allow us to resolve the receptive fields of HSS cells. Therefore, only HSN and HSE vector fields were considered for detailed analysis (Figure 5A, B), where the arrow’s length and direction represent the response’s relative local strength and preferred direction, respectively.

**Figure 5.**
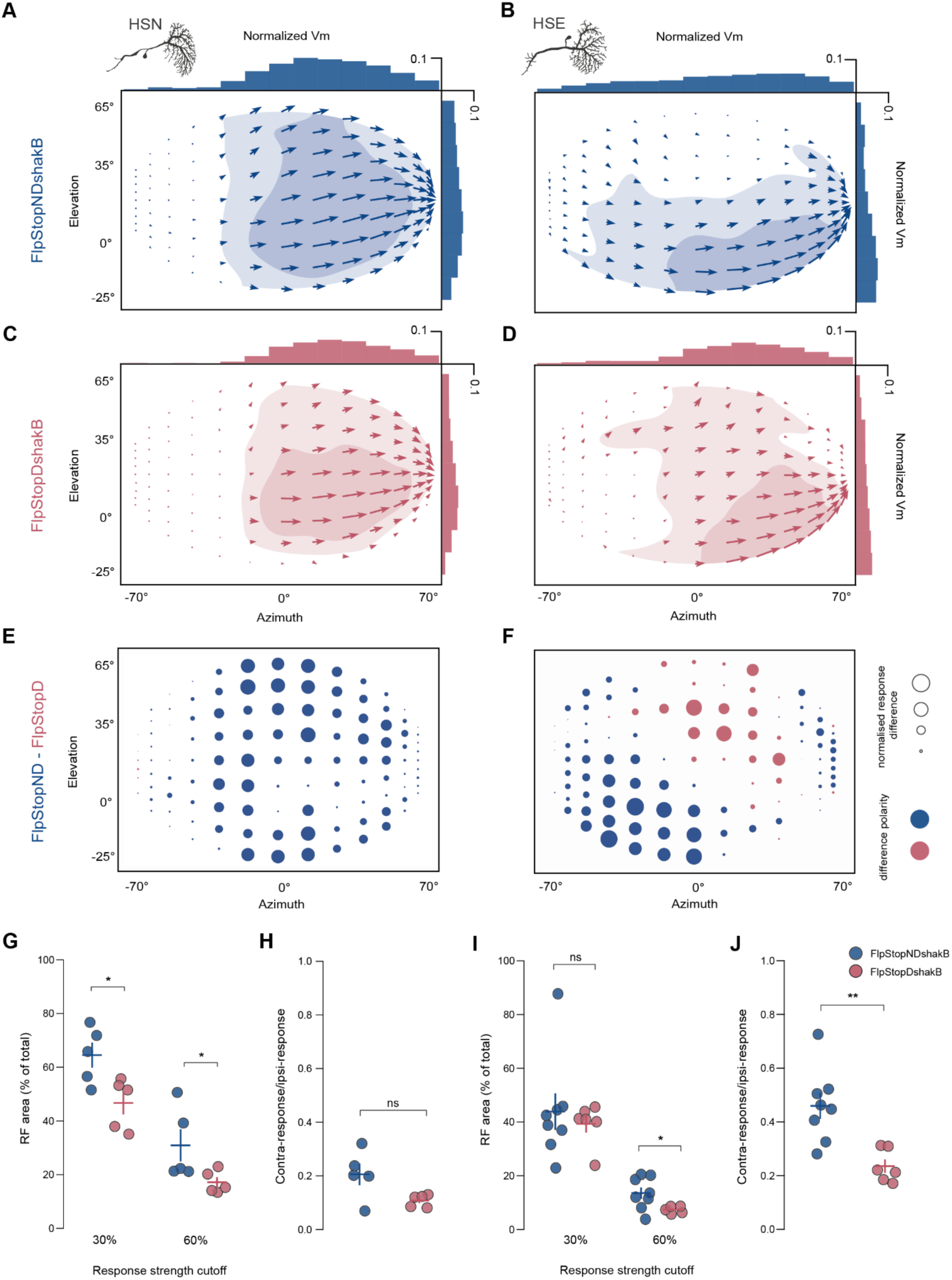
Gap junctions shape receptive field structure. (A-D) Spatial receptive fields reconstructed from the responses to local motion stimulus (see methods) for HSN (A, C) and HSE (B, D) cells in FlpND and FlpD flies. Light- and dark-shaded areas represent 30% and 60% of the maximal strength of the response for each cell type. (E-F) Differences in the amplitude of local motion responses in HSN and HSE cells between FlpND and FlpD flies. The size of the circle depicts the magnitude of the response difference, and the color represents polarity. Blue depicts a positive and pink a negative difference. (G) Size of the receptive field, corresponding to the shaded regions in (A, C) for individual HSN cells. Bars: mean ± SEM. (H) Relative strength between the contralateral and ipsilateral visual fields of individual HSN cells. Bars: mean ± SEM. (I) Same as in (G) for HSE cells. (J) Same as in (H) for HSE cells. (G, I) one-sided Mann-Whitney U test, (H, J) two-sided Mann-Whitney U test. ∗*p* < 0.05, ∗∗p < 0.01. The number of recorded cells and exact *p*-values for each experiment are listed in Suppl. Table1.

As shown previously^5,28^ and in correspondence with the dendritic arborizations, RFs of HS cells in FlpND flies are aligned horizontally, with their RF centers differing in elevation - HSN being higher than HSE. Both types of HS cells showed sensitivity to local back-to-front motion in the contralateral visual field, comprising 25.3% of the total sensitivity of HSN cells and 36.6% of HSE cells (Figures 5A,B, S8E,F). The motion sensitivity of HSE cells was detected along the entire span of measured azimuth (−70° to 70°). This complex structure of local motion sensitivity on the contralateral side is inherited from contralateral elements connected to HS cells.

In FlpD mutant flies, the size of the spatial receptive fields of both HSN and HSE cells was largely reduced (Figures 5C, 5D, 5G, 5I, Figure S8G, S8H). While strong responses to ipsilateral horizontal motion along the equator were preserved in both HSN and HSE cells, HSN cells showed reduced responses in the fronto-dorsal and ventral areas, and HSE cells lost sensitivity to the motion in the contralateral field almost entirely. These changes suggest that motion sensitivity in these areas is a result of lateral interactions of HS with other tangential cells: likely horizontal-motion-sensitive ipsilateral neurons for HSN cells, e.g. CH-cells, and contralateral horizontal-motion-sensitive neurons for HSE cells, e.g. H2. Interestingly, direction-selective responses of HSE cells in mutant flies were enhanced in the fronto-dorsal area, indicating a potential role of gap junction-mediated inhibitory inputs in shaping the RF of these cells^29,30^. The described differences in RFs arise mainly through changes in inputs to their preferred direction (PD), with a complete abolition of contralateral responses in HSE neurons (Figures S8A-D).

Altogether, the shape of HS cell RF is strongly modulated by lateral interactions via gap junctions. In particular, HSE neurons acquire back-to-front motion sensitivity in the contralateral area through a direct electrical coupling with contralateral horizontal-motion-sensitive neurons, enhancing responses to yaw-rotation tuning as suggested previously in blowflies^13^.

### Dye coupling unveils the electrical connectivity network of HS cells

To visualize electrical coupling, we filled individual HS cells by injecting neurobiotin, a molecule that can diffuse through gap junctions. Subsequent visualization of neurobiotin using fluorescently tagged streptavidin revealed that HS cells in FlpND flies are electrically coupled to each other as well as to postsynaptic interneurons, motoneurons, and descending neurons (Figure 6A, 6B), as shown in numerous previous studies^5,16, 31–34^. HSN cells are strongly connected with two descending neurons, including DNp15 (DNHS1). Similar to blow flies, we also observed gap junctions between HS cells and neck motoneurons. Specifically, HSN and HSE form electrical connections with VCNM-like neurons (Ventral Cervical Nerve Motor Neuron), and HSS cells form electrical synapses with CNM-like neurons (Cervical Nerve Motor Neuron). Unlike wild-type flies, FlpD mutant flies showed little to no dye coupling between ipsilateral HS cells, indicating that axo-axonal gap junctions between HS neurons are absent or severely perturbed (Figures 6D, 6E). Additionally, coupling between HS cells and another class of ipsilateral LPTCs, presumably CH cells, was largely abolished in FlpD mutant flies.

**Figure 6.**
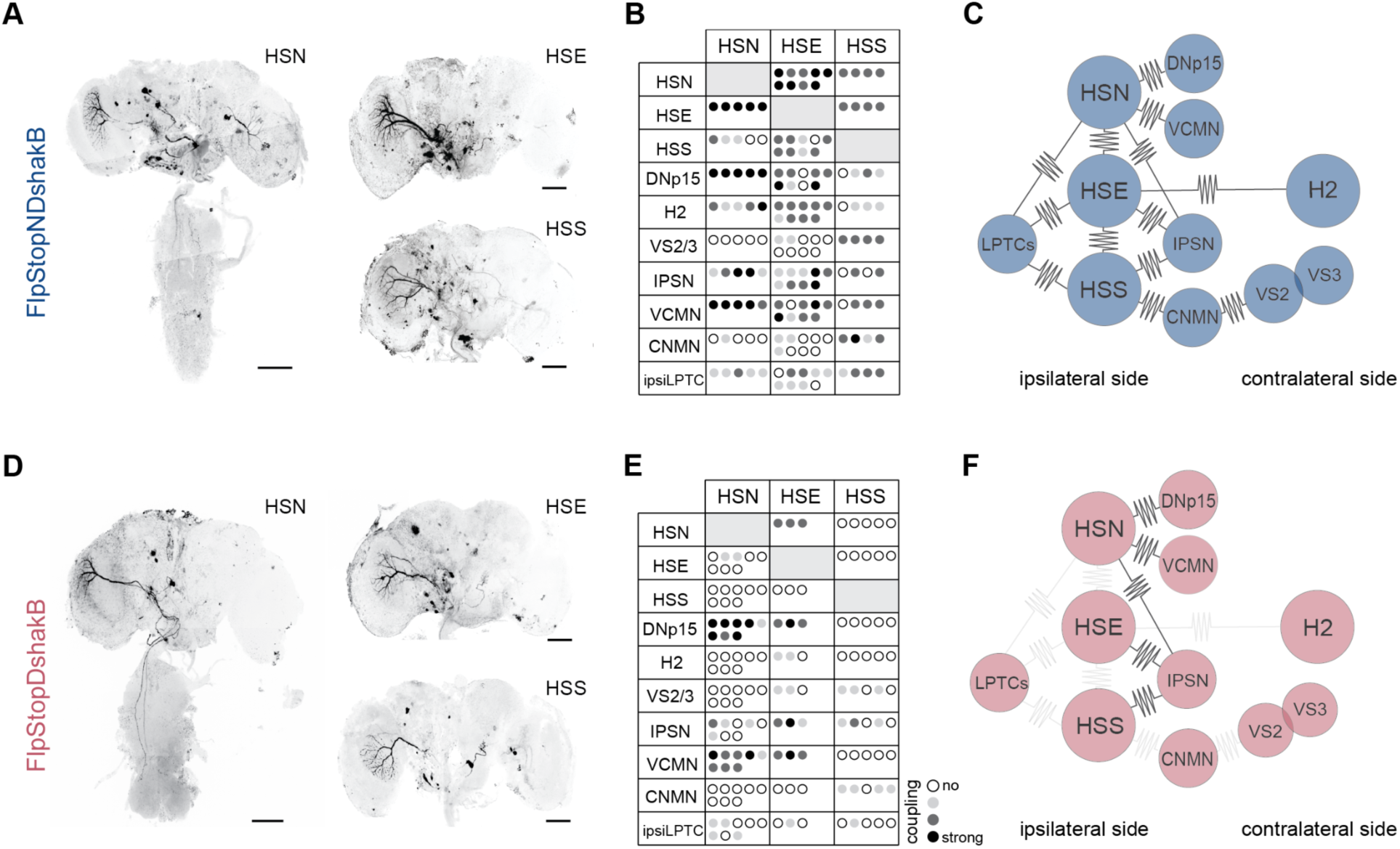
Specific sensory-to-sensory coupling disruption, not sensory-to-motor, in FlpD mutant. (A) Examples of neurobiotin injections into individual HS cells in FlpND flies. (B) Quantification of electrical coupling from individual HS cell dye fillings. The table summarizes the most prominent electrical partners. Each dot represents an individual experiment. Dot opaqueness represents coupling strength inferred from the staining intensity. (C) Diagram summarizing electrical partners of HS cells in FlpND. (D) Same as in (A), but for FlpD flies. (E) Same as in (B), but for FlpD flies. (F) Same as in (C), but for FlpD flies. (A, D) Scale bars: 50 μm.

While connections between LPTCs in FlpD mutant flies were abolished, electrical coupling between HS cells and postsynaptic descending neurons was largely preserved. This suggests that gap junctions formed between HS cells and postsynaptic neurons differ molecularly from those formed between LPTCs, and are likely composed of either undisrupted *shakB* isoforms or innexins other than *shakB*. Interestingly, unlike FlpD, HS cells in *shakB*^2^ mutant flies do not show any dye coupling^16^ despite expressing an additional isoform (*shakB-RF*) (Figure S3B).

Apart from ipsilateral electrical coupling, HS cells also formed extensive connections with neurons in the contralateral hemisphere. All three HS neurons were coupled to bilateral neurons in the inferior posterior slope (IPSN) that form a bridge between the tangential cells of the two hemispheres. This coupling is maintained in FlpD mutant flies. We also observed dye coupling between HSS and contralateral VS2 and VS3 neurons mediated via CNMNs. HSE cells form electrical synapses with contralateral tangential cells H2. H2-HSE connection is substantially weakened in FlpD mutant flies and, therefore, can explain the difference in the structure of HSE receptive fields observed in FlpND and FlpD flies.

Receptive fields of HSE cells in DB331xFlpND flies did not show differences from wild-type flies (Figures S8 I-N). This result is in line with the pattern of dye coupling (Figures S8K,S8M), showing that the gap junctions in HS cells are largely unaffected after cell-specific inactivation of *shakB*, despite a reduction of protein amount in LPTC terminals (Figure 3F). However, as mentioned before, cell-specific inactivation worked completely on rare occasions in VS cells (Figure S7C), indicating that *shakB* can be disrupted in a cell-specific manner with GAL4 lines that express in early development.

Overall, the neurobiotin injections into HS cells revealed a significant reduction in the strength of their electrical coupling in FlpD mutant flies (Figure 6C,F). However, this reduction was limited to connections between LPTCs, while connection between HS cells and postsynaptic interneurons, descending neurons, and motoneurons remained intact, making FlpD mutant a favorable model for studying the behavioral role of gap junctions. Crucially, the reduction of contralateral electrical connectivity enables us to test the involvement of gap junctions in integrating binocular motion cues.

### Gap junctions arbitrate binocular behavioral instructions

To study the behavioral role of gap junction-mediated binocular interactions, we compared the turning response of FlpD and FlpND flies to full-field and unilateral FtB and BtF rotation. Similar to wild-type flies, FlpND flies turned in the direction of unilateral BtF and full-field rotation, and against the direction of unilateral FtB motion, demonstrating that the insertion of the genetic cassette does not affect the turning responses of the fly (Figure 1D, 7A). However, the overall responses were stronger for FlpND flies in comparison to CantonS flies, which can be attributed to the differences in the genetic backgrounds of these two lines. The relative contribution of smooth and saccadic turning in FlpND was similar to that observed for wild-type flies (Figure 1Hi, 7Ai *bottom*). Similarly, FlpD flies exhibited a strong turning response in the direction of full-field rotation (Figure 7Ci), showing that the optomotor response is remarkably robust to perturbations in the LPTC network, in line with afore-described observations in flies with silenced HS and VS neurons (Figure 2C). Likewise, FlpD flies turned toward BtF motion (Figure 7Cii). However, in contrast to FlpND flies that turned in a saccadic manner against FtB stimuli, FlpD flies inverted the behavioral strategy and response direction, turning on average smoothly with the direction of the stimulus (Figure 7Ciii).

**Figure 7.**
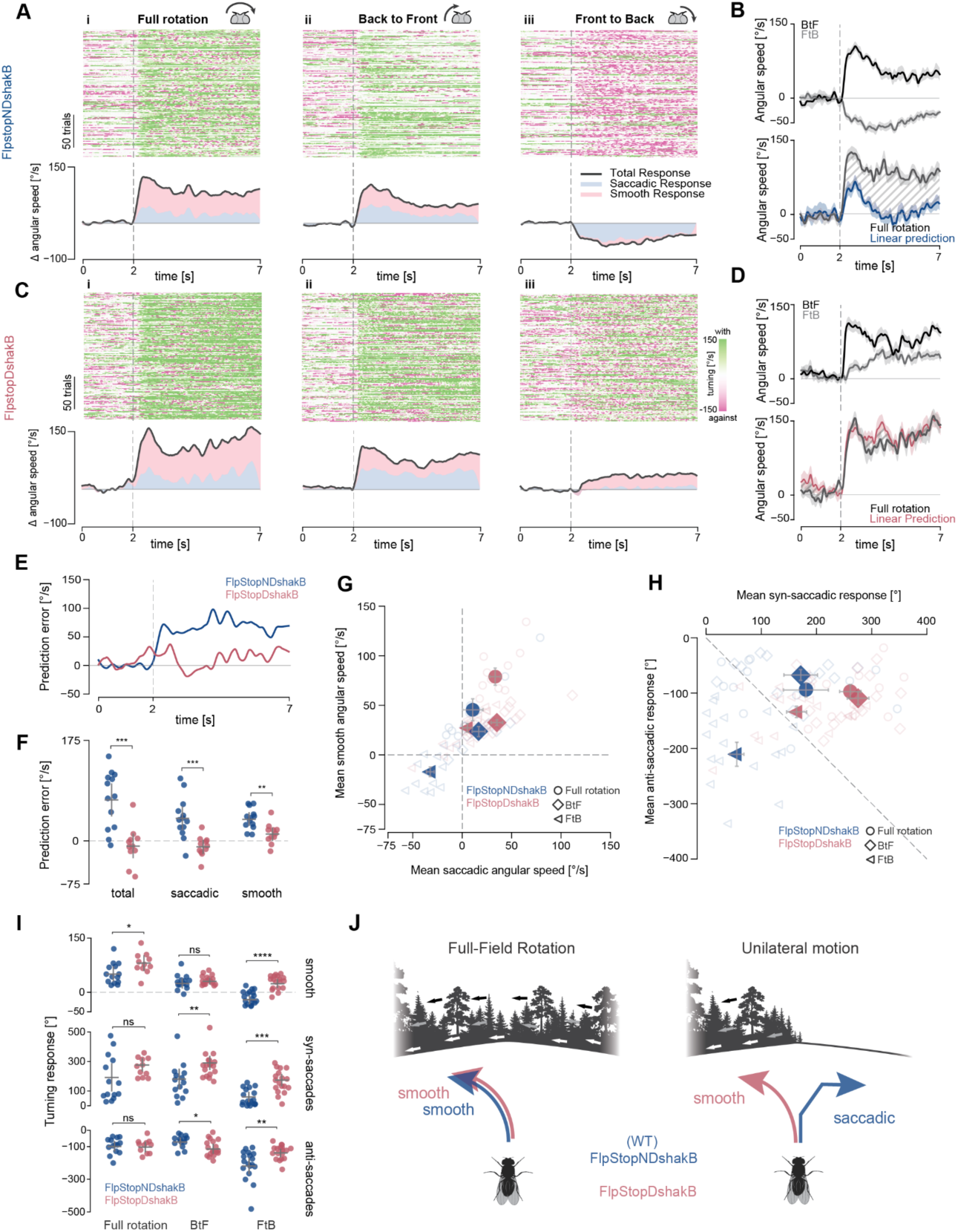
Binocular control of walking behavior requires gap junction connectivity. (A) Turning response of FlpND and FlpD flies to (i) full-field, (ii) unilateral Back to Front (BtF) and (iii) unilateral Front to Back (FtB) rotation. *Top.* Angular speed raster plots showing 200 randomly selected trials. Each row corresponds to one trial, each column to one frame (∼16 ms). *Bottom.* Stacked plot showing mean angular speed and the contribution of smooth and saccadic turning. (B) *Top.* Angular speed of FlpND flies in response to FtB and BtF rotation (mean ± SEM). *Bottom.* Comparison of the predicted and the actual mean angular speed in response to full-field rotation for FlpND flies. Shaded region shows the difference between predicted and actual response. (C) Same as in (A) but for FlpD flies. (D) Same as (B) but for FlpD flies. (E) Time series of mean prediction error. (F) Mean prediction error across the trial period per fly. Bars: mean ± SEM. (G) Summary plot of the mean of smooth and saccadic angular speed across experiments. Empty and filled markers show the mean per fly and mean ± SEM for each genotype, respectively. (H) Summary plot of the mean of syn-saccadic and anti-saccadic turns made. Empty and filled markers show the mean per fly and mean ± SEM for each genotype, respectively. (I) Mean of smooth (top), syn-saccadic (middle) and anti-saccadic (bottom) turns made per trial. Bars: mean ± SEM. (J) Schematic summary. Contralateral HSE to H2 electrical connectivity arbitrates the change between optomotor and saccadic counter-response. Disrupting the contralateral connectivity disrupts this finely tuned bilateral course control, disrupting the counter-saccadic response. (F, I) two-sided Mann-Whitney U test was applied, ∗*p* < 0.05, ∗∗p < 0.01, ∗∗∗p<0.001, ∗∗∗∗p<0.0001. No asterisk: not significant. The number of flies per genotype and exact *p*-values for each experiment are listed in Suppl. Table1.

Consistent with wild-type flies, the optomotor response to full-field rotation exhibited significant differences from the linear prediction in FlpND flies (Figures 7B,7E,7F). However, in FlpD flies, the optomotor response closely matched the linear prediction (Figures 7D-F), indicating that binocular interactions facilitated by gap junctions are directly responsible for the non-linear summation of motion cues from both halves of the visual field. Importantly, the linearization of the optomotor response is primarily attributed to changes in smooth and saccadic responses, as evidenced by the reduction in prediction error for both types of movements (Figure 7F). Notably, in FlpD flies subjected to front-to-back (FtB) rotation, smooth turning in the direction of the stimulus was enhanced, while saccadic counter turns were diminished (Figures 7G-I, S9B).

In summary, the abolition of electrical coupling has a profound impact on binocular behavioral instructions. It enhances smooth and saccadic syn-directional turning while significantly reducing saccadic counter turns. As a result, the fly’s binocular integration undergoes a shift in complexity. It transitions from non-linearly integrating monocular information for guiding course corrections to linearly summing binocular cues, constraining the behavioral repertoire of the fly (Figure 7J).

## Discussion

The intricate wiring of the nervous system relies on a variety of cellular mechanisms to facilitate communication between neurons. One such mechanism is electrical coupling by gap junctions, specialized channels between adjacent cells. Their computational roles have been proposed to contribute to the synchronization or desynchronization of network activity^35–37^, improve sensory estimates of the environment^18^, act as computational switches in sensory processing^38^, and be relevant for cell-intrinsic stability^16^. However, the implications of such computations for behavior remain largely unknown. Here we present evidence that specific gap junction connectivity plays a decisive role in sensorimotor transformations. First, we uncovered the richness and precision of optic-flow-based navigation by expanding the classical optomotor paradigm (Figure 1A-D). We show that flies change their locomotion strategy depending on the statistics of the visual stimulus, changing the nature and direction of their turning responses (Figure 1D-F). These changes underlie a non-linear visuomotor transformation since the summation of each monocular stimulus can’t account for the binocular reaction (Fig. 1G-I). This computation is mediated partly by a subset of LPTCs (Figure 2), namely HS-cells, known to be connected via gap junctions contralaterally to H2^13^. The *shakB* gene has been identified as a key player in forming these gap junction channels^16^ (Figure 3), and several studies have implicated electrical connectivity in the LPTC network with improved efficiency of optic-flow-based computations^13,18,19,39,40^. Yet, these studies remain beyond experimental verification. By establishing and thoroughly characterizing a new *shakB* mutant line (Figures 3, 4, 5, 6, S3, S4, S5, S6, S7 & S8), we could demonstrate that the behavioral deficits observed by silencing LPTCs (Figure 2) are due to the loss of interhemispheric gap junctional connections between HS and H2 cells (Figures 5, 6, 7). These results expand the computational roles mediated by gap junctions and implicate them with non-linear operations with a decisive role in animal course control.

### A repertoire of course control behaviors

Flies’ remarkable course control behaviors have been postulated to reside largely in the algorithms embedded in the connectivity among LPTCs, the so-called “cockpit of the fly”^3^. Yet, the relevance of the connectivity between the LPTCs is poorly understood, partly because the behavioral repertoire has been restricted mainly to the classical optomotor response - a reflexive response that does not require the complexity seen in these circuits. Here, by employing close-loop freely walking behavioral assays (Figure 1A-C), we have identified strikingly different visuomotor behaviors, opening the path for a dissection of the underlying neuronal mechanisms. Specifically, we showed that the turning direction and behavioral strategy (saccadic or smooth) depend on the optic flow’s global properties. These results, at first glance, appear counterintuitive because previous studies have shown that syn-saccades are triggered when smooth turning is insufficient to counteract retinal slip^41^. But anti-saccadic responses, as seen during FtB motion, would increase retinal slip. Clearly, the fly extracts different behavioral meanings from distinct optic flow patterns that go beyond course stabilization to aid visual navigation across natural environments (Figure 7J).

Our observations differ from recent reports of anti-directional, saccade-independent turning in response to long-lasting, high-contrast visual stimuli^21^. While we did observe a few anti-directional responses to full-field rotation in our experiments, they were mainly saccadic. Yet, the average response of the fly was in the direction of motion across the entire trial length, composed of smooth turns and syn-saccades. Strong and immediate anti-saccadic reactions were observed specifically for FtB motion. This difference may arise due to our close-loop configuration, suggesting that the gain of visual feedback affects the turning response of the fly. Finally, unlike other studies^42^, we present the visual stimulus from above and mostly target the dorsal visual field. Recent work has shown that moths have different flight stabilization strategies depending on whether the optic flows are presented from above or below^43^, consistent with changes in natural image statistics across elevation. Adaptation of such panoramic natural image statistics has been recently shown to occur in the mammalian retina as well^44^, indicating that these adaptations are a general phenomenon of visual systems. Thus, the panoramic position of optic flow likely plays a crucial role in generating behavioral instructions.

### Molecular Diversity of Gap Junction Channels

The *shakB* gene in *Drosophila* produces multiple isoforms (Figure 3B, S3B), resulting in distinct protein variants. These isoforms exhibit specific temporal and spatial expression patterns, suggesting their involvement in the formation of different types of gap junction channels. However, the function of this molecular diversity remains poorly understood. This becomes evident when observing the phenotypic discrepancies between the FlpD and *shakB*^2^ mutant flies. We observed systematic variations in dye-coupling, hyperpolarization patterns, and the strength of responses in different neuron types. Most strikingly, whereas HS cells in *shakB*^2^ mutant flies exhibit dramatically perturbed visual responses and a complete absence of dye coupling^16^, HS cells in FlpD flies largely preserve their visual response properties (Figure 4) and show very specific changes in their electrical coupling (Figure 6). The potential sources of these discrepancies include (i) a reduced penetrance of the FlpStopDshakB allele, which was evident in female flies, probably because *shakB* is located on the X chromosome. We circumvented this problem by performing all our experiments on male flies. (ii) Additional mutations accumulated or generated during EMS-induced mutagenesis in the *shakB*^2^ mutant, or (iii), putatively dominant negative effects of the additional *shakB-RF* isoform that remains intact and is expressed at a relatively high level throughout the development and during adult stage (Figure S3B). Given the difficulties in ruling out the underlying differences in the *shakB*^2^ mutant, new genetic approaches similar to our FlpStop approach are needed to understand the specific computational implications of gap junctions.

### Electrical connectivity logic

Why would the disruption of one innexin gene differentially affect electrical synapses formed by HS cells? Given that the shakB gene produces multiple isoforms, the most obvious explanation is that HS cells utilize different protein variants to form gap junctions with distinct synaptic partners. Several lines of evidence corroborate this hypothesis. Different isoforms of the *shakB* form various channels with distinct properties that shape a unique current flow pattern in a given neural circuit^45^. For example, heterotypic channels were demonstrated to form rectifying electrical synapses in the giant fiber system, while homotypic channels exhibit symmetrical voltage responses^45^. Specifically, the giant fiber neuron was shown to express the ShakB(N+16) to form gap junction channels with postsynaptic motoneurons that express the ShakB(Lethal) variant. In these experiments, only ShakB(N+16) expression and not ShakB(N) rescued the connectivity phenotype in the giant fiber system in a *shakB*^2^ background^45^, suggesting the underlying connectivity logic might reflect differences in the functional connectivity properties too. Thus, coupling between LPTCs and downstream DNs and motor neurons could be composed of rectifying electrical synapses. A thorough genetic and physiological analysis would be required to unravel the exact molecular composition and physiological properties of ShakB electrical synapses in tangential cells.

### Disrupting *shakB*

Previous cell-specific knock-down attempts by RNAi^16^ and our attempts with CRISPR/Cas9 (data not shown) did not significantly reduce the electrical coupling formed by LPTCs. These technical challenges are rather puzzling, considering that the predicted lifespan of gap junction proteins in *Drosophila* is several hours to several days^46,47^. However, the analysis of several driver lines for LPTC-specific inversion of the FlpStop cassette shows that this prediction may not be accurate for ShakB protein in all cells. The induction of gene disruption in LPTCs at around developmental stage P9 results in a stronger reduction of protein signal in adult animals than induction at later stage P12 (Figure 3E, 3F). Despite strong reduction in the protein amount induced by early and unspecific DB331 line, we observed only on rare occasions a full disruption of gap junctions in VS cells (Figure S7C). This effect is likely due to an earlier onset of GAL4 in these cells. Importantly, a previous study has shown that a weak pan-neuronal expression of Shakb(N+16) can rescue electrical connectivity in the giant fiber system^45^, or in our case, a severe reduction in the amount of ShakB did not result in a loss of electrical synapses formed by LPTCs. This shows that even a small amount of protein can form functional gap junction channels. Thus, future work aimed at disrupting electrical synapses in a cell-specific manner will also need to consider the exact onset of the driver lines during development.

### Visual response properties in the absence of ShakB

As previously reported for *shakB*^2^ flies, we also observed β-oscillations in the resting membrane potential in FlpD mutant flies^16^ (Figure 4F, 4G). Interestingly, we show that these changes are cell-intrinsic by correlating the oscillations of membrane potential with the absence of dye coupling after LPTC-specific disruption of *shakB* (Figure S7C), indicating that gap junctions formed between HS cells facilitate the dissipation of cell-intrinsic noise that otherwise can interfere with signals arriving from presynaptic cells. Contrary to a previous report on *shakB*^2^ mutant flies^16^, the main direction-selective properties, as well as the velocity and contrast tuning, are not affected in FlpD mutant flies (Figure 4K-4M). The only observed difference was an increase in hyperpolarization amplitude in response to gratings moving in the null direction for ON edges (Figure 4 I-M), indicating that this is not a cell-extrinsic phenomenon. The most striking effect is observed in the structure of the receptive fields of HS cells. For example, in wild-type animals, HSE cells receive strong contralateral excitatory input for motion in the preferred direction. This input was absent in FlpD mutant flies (Figure 5, S8A-D). In line with previous work^13^ and intracellular dye fillings described above, this contralateral input arrives from back-to-front-motion-sensitive H2 neurons. This binocular connectivity has been proposed to be required for accurate interpretation of optic flows dominated by horizontal components, such as yaw rotation and translation^13^. Therefore, the loss of the HSE-to-H2 connection would restrict the analysis of the optic flow to individual hemispheres, limiting the ability of the fly to differentiate global optic flow patterns and instruct adequate steering behaviors (Figure 7J).

### Course control in the absence of ShakB

Because different algorithms can solve the same computational problem, a rigorous characterization of the stimulus–behavior mapping is required to understand their neuronal implementation^48^. Thus, we performed a behavioral analysis comparing two sets of FlpStop flies: one with the cassette in the disruptive orientation and the other in the non-disruptive orientation. These flies share an identical genetic background, and any behavioral discrepancies can be attributed solely to the disruption of the *shakB* gene. Remarkably, we observed a similar behavioral phenotype to that observed in our silencing experiments targeting LPTCs (Figure 2), suggesting that the mechanism behind bilateral visual control relies on the electrical coupling between these neurons. The extent of behavioral disruption in FlpD mutants appeared to be more pronounced compared to our silencing experiments, where we observed an initial counter-saccadic turn in response to FtB stimuli (Figure 2B). This discrepancy is likely attributable to the silencing approach using Kir2.1, which depends on expression levels. In HS cells, this silencing approach has been shown to leave a residual direction-selective activity^10^. Interestingly, in both cases, the linear sum of the behavioral response to monocular stimuli matches the binocular optomotor response (Figures 2C, 7D-E), an equivalence that is absent in wild-type and FlpND flies due to a striking non-linear interaction between two hemispheres (Figures 1G, 7B, 7E). Consequently, our results provide compelling evidence implicating a specific LPTC network computation in behavior. These findings underscore the computational and behavioral importance of gap junctions within the insect nervous system, connections that have been predominantly overlooked in connectomic analysis due to the challenges associated with their visualization. Moreover, they suggest that gap junctions can facilitate non-linear operations that play a decisive role in animal course control rather than merely averaging neighboring signals. However, a significant question remains unanswered. What are the underlying circuit motifs and biophysical implementations underlying this non-linear, gap-junction-mediated binocular interaction? Although our research points to the relevance of the interaction between HSE and H2, it is important to consider that this computation occurs within a circuit context that necessarily involves additional components. For example, saccadic turns are governed by defined pathways that project symmetrically to the ventral nerve cord and are believed to suppress each other reciprocally^49^. More than 50 years ago, the optomotor response^1^ laid the foundation for subsequent research on the neuronal implementations of motion vision^2^. Likewise, we believe that the refinement of course control behaviors, as done in this study, is a requirement for a comprehensive understanding of the neuronal mechanisms of vision in flies^48,50^.

## Acknowledgment

We thank Georg Ammer and Alexander Borst for sharing anti-ShakB serum antibodies. We thank Nélia Varela and Eugenia Chiappe for the *w1118;+;10XUAS-IVS-eGFPKir2.1/TM6B* fly line, Augustin Hrvoje for the *shakB*^2^ line, as well as Jesse Isaacman-Beck and Thomas R Clandinin for the gift of *y*^1^*,w*;20XUAS-IVS-PhiC31;+* fly line. We also thank Armel Nicolas and Tomas Masson for the proteomic analysis, Ece Sönmez for help with fly crosses and dissections for protein analysis, and Lisa Hofer for assistance with the reconstruction experiments. We are particularly grateful to members of the Siekhaus, the Kondrashov, and the Chiappe group for providing material support and technical advice. We are grateful to Daria Siekhaus, Eugenia Chiappe, Ben deBivort, and all the members of the Joesch laboratory for valuable discussions and comments on the manuscript. Stocks from the Bloomington Drosophila Stock Center (NIH P40OD018537) and the Vienna Drosophila Resource Center were used in this study. The Scientific Service Units of ISTA supported the project through resources provided by the Imaging and Optics Facility, MIBA Machine Shop, and the Lab Support Facility, as well as Vienna *Drosophila* Research Centre. This work was funded by the Deutsche Forschungsgemeinschaft (DFG, German Research Foundation) as part of the SPP 2205 - 429960716.

## Author contributions

V.O.P., R.S., and M.J. designed the study. V.O.P. performed the electrophysiological experiments, intracellular dye fillings, neuronal tracing and reconstructions, protein analysis, immunochemistry, acquisition and analysis of microscopy images. O.S., R.S., and V.O.P. performed the analysis of electrophysiological data. O.S. and M.J. built the electrophysiology setup. R.S. built and designed the behavioral setup, and performed all behavioral experiments and analyses. V.O.P., R.S., and M.J. wrote the manuscript.

## Materials and Methods

### Fly stock list

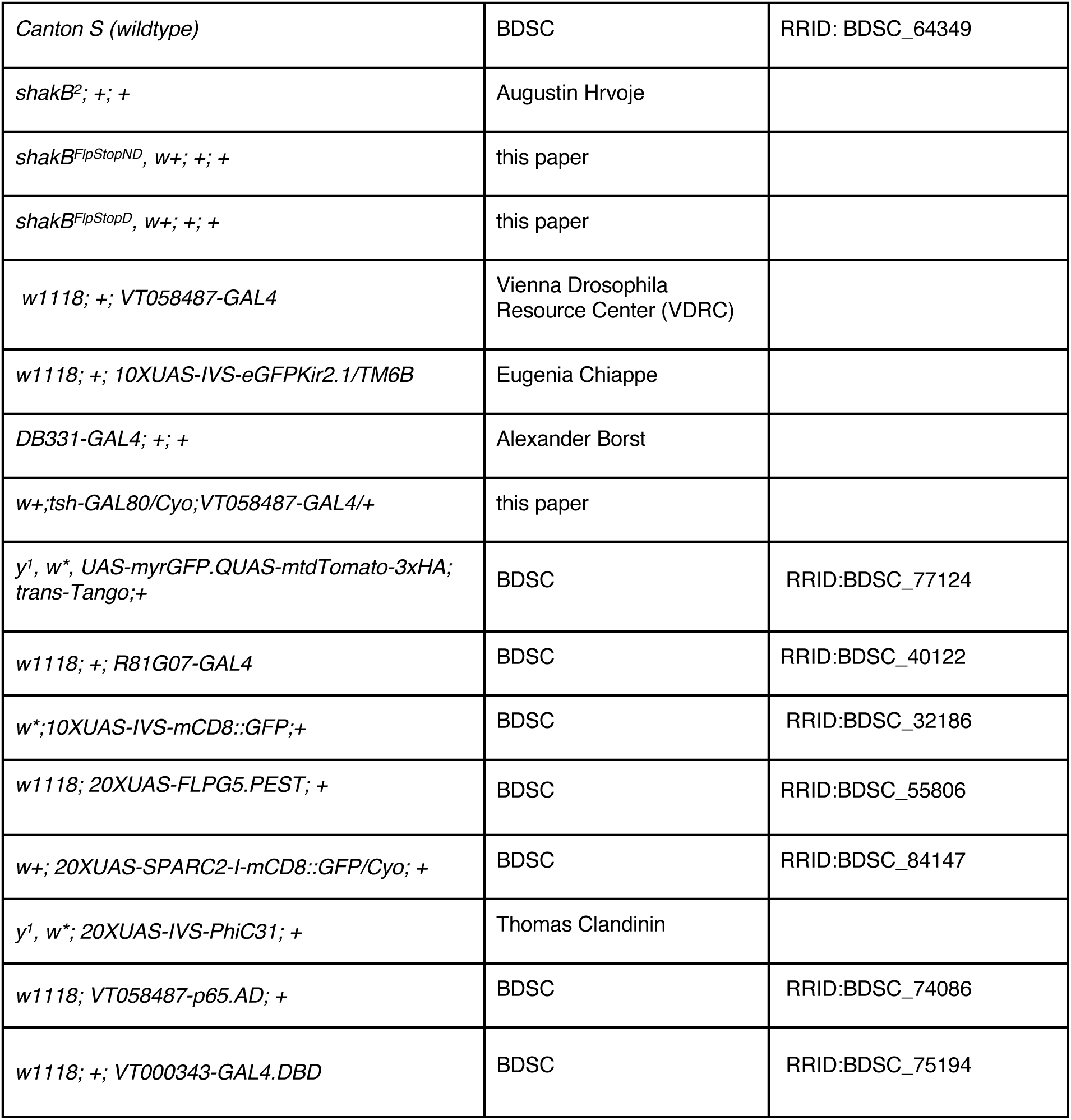

### Detailed fly genotypes used

Figure 1B-1L, S7A, S7B

*Canton-S*

Figure 2A-2C, 2F-2J

*w+; tsh-GAL80/+;VT058487-GAL4/10XUAS-IVS-eGFPKir2.1*

Figure 2D-2J

*w+; +;10XUAS-IVS-eGFPKir2.1/+*

Figure 3C, 3E

*shakB*^2^*; 10XUAS-IVS-mCD8::GFP/+; VT058487-GAL4/+*

*shakB^FlpStopND^; 10XUAS-IVS-mCD8::GFP/+; VT058487-GAL4/+*

*shakB^FlpStopD^; 10XUAS-IVS-mCD8::GFP/+; VT058487-GAL4/+*

Figure 3D, S3A, S3B,

*shakB*^2^*; +; +*

*shakB^FlpStopND^, w+; +; +*

*shakB^FlpStopD^, w+; +; +*

Figure 3E, 3F, S3C

*shakB^FlpStopND,^ DB331-GAL4; 20XUAS-FLPG5.PEST/+; +*

*shakB^FlpStopND^, w+; 20XUAS-FLPG5.PEST/+; VT058487-GAL4*

Figure S5

*w1118; 20XUAS-SPARC2-I-mCD8::GFP/VT058487-p65.AD, 20XUAS-IVS-PhiC31; VT000343-GAL4.DBD/+*

*shakB*^2^*; 20XUAS-SPARC2-I-mCD8::GFP/VT058487-p65.AD, 20XUAS-IVS-PhiC31; VT000343-GAL4.DBD/+*

*shakB^FlpStopD^; 20XUAS-SPARC2-I-mCD8::GFP/VT058487-p65.AD, 20XUAS-IVS-PhiC31; VT000343-GAL4.DBD/+*

Figure S6

*w1118, UAS-myrGFP.QUAS-mtdTomato-3xHA; trans-Tango; R81G07-GAL4/+*

*shakB*^2^ *UAS-myrGFP.QUAS-mtdTomato-3xHA; trans-Tango; R81G07-GAL4/+*

*shakB^FlpStopD^ UAS-myrGFP.QUAS-mtdTomato-3xHA; trans-Tango; R81G07-GAL4/+*

Figure 4B-4M, S7D-S7I

*shakB*^2^*; +; +*

*shakB^FlpStopND^, w+; +; +*

*shakB^FlpStopD^, w+; +; +*

*shakB^FlpStopND^, DB331-GAL4; 20XUAS-FLPG5.PEST/+; +*

Figure 5A-5J, 6A-6F, 7A-7K, S8A-S8H, S9A-S9E

*shakB^FlpStopND^, w+; +; +*

*shakB^FlpStopD^, w+; +; +*

Figure S7C, S8I-S8N

*shakB^FlpStopND^, DB331-GAL4; 20XUAS-FLPG5.PEST/+; +*

### Fly husbandry

*Drosophila melanogaster* were reared on a standard cornmeal-molasses agar medium at 18°C and 60% humidity and kept on a 12h light/12h dark cycle. Experiments were performed with 2-to-6-day old flies. Due to reduced penetrance and high phenotypic variability in female mutant flies, we used only male flies.

### Generation of shakB^FlpStop^ transgenic flies

pFlpStop-attB-UAS-2.1-tdTom (addgene #88910) donor plasmid was injected into shakB[*MI15228*] flies together with φC31 integrase-expressing transgene (BestGene Inc.). The orientation of the FlpStop cassette was identified using primers MiL-F, FRTspacer_5p_rev, and FRTspacer_3p_for from^23^.

### Protein isolation and quantification of ShakB

To extract insoluble protein fraction, approximately 150 brains of 3-to-5-day-old male flies were homogenized in 300 μl of extraction buffer (20 mM Tris pH 7.6, 50 mM NaCl, 1 % Triton X-100, 1XHalt Protease inhibitor Cocktail (Thermo Fisher, 78429)), and incubated for 30 min on ice. Homogenates were centrifuged for 60 min at 15 x 1000 g in 4 °C. Supernatant was discarded, and the remaining pellet was used for the isolation of insoluble proteins. For that, the pellet was resuspended in the SDS extraction buffer (50 mM Tris pH 7.6, 5 mM EDTA, 4 % SDS), and incubated at 95 °C for 10 min. Supernatants were collected after centrifugation for 10 min at 15 x 1000 g at room temperature and were used for protein quantification with BCA protein assay (Thermo Fisher, 23225).

For western blot analysis, 10 μg of each protein sample were mixed with Laemmli’s buffer, boiled for 5 min, and subjected to SDS-PAGE using 4–20 % TGX Stain-Free Precast Gels (Bio-Rad, 4568095). Gels were activated by UV exposure for 2 min using a Bio-Rad Chemidoc MP imager. Proteins were transferred to LF PVDF membranes (Bio-Rad, 1620263) using a Transblot Turbo apparatus (Bio-Rad). The membrane was incubated in EveryBlot Blocking Buffer (Bio-Rad, 12010020) and immunoblotted following standard protocols.

The following antibodies were used for the western blot analysis: anti-ShakB (1:3000, Innovagen AB), IRDye 800CW goat anti-rabbit (1:15000, LI-COR Biosciences). Total lane signal was detected using Stain Free Blot application and ShakB signal was detected using IRDye 800 CW application (Bio-Rad Chemidoc MP imager).

The immunoblots were repeated three times. ShakB signal was quantified and normalized to the total lane signal using Imagelab 4.1 (Bio-Rad).

### Proteomic analysis

For protein sample preparation, brains of 2-5 days old flies were dissected. All samples (3 genotypes, 4 replicates; 10 or 6 pooled dissected fly brains per sample for replicate 1 and replicates 2, 3, and 4, respectively) were processed in 4 replicate-specific batches with the iST-NHS kit from PreOmics GmbH using the standard manufacturer’s protocol with the following modifications: samples were lysed in 100 µL iST-LYSE buffer, boiled at 95 °C for 5 min, then sonicated for 10 cycles of 30 s each on/off in a Bioruptor plus (Diagenode) in presence of 50 mg Protein Extraction beads (C20000021, Diagenode); samples were trypsin digested for 2 h 30 min, then labeled with TMT-6plex (ThermoScientific, lot # VI307213) according to the manufacturer’s instructions (one bridging channel, consisting of a mix of all 20 samples, was included in each combined TMT sample). Combined TMT samples were dried, re-dissolved in 45 µL 100 mM NH4OH, then loaded onto an ACQUITY UPLC BEH C18 column (130 Å, 1.7 µm, 2.1 mm x 150 mm, Waters) on an Ultimate 3000 UHPLC (Dionex) and fractionated into 24 fractions by High pH Reversed Phase chromatography (solvent A: deionized water + 10 mM NH4OH; B: 90 % LC-grade Acetonitrile + 10 mM NH4OH; flow: 0.15 ml/min; gradient: 0-4 min = 1 % B, 115 min = 25 %, 140 min = 40 %, 148-160 min = 75 % followed by re-equilibration at 1 % B). Fractions were combined at mid-gradient, re-dissolved in 50 µL iST-LOAD and sent for MS analysis.

All samples were analyzed by LC-MS/MS on an Ultimate 3000 nano-HPLC (Dionex) coupled with a Q-Exactive HF (ThermoFisher Scientific). Spectral data were acquired on data-Dependent Acquisition (Full MS/dd-MS2); chrom. peak width (FWHM) 20 s, MS1: 1 microscan, 120,000 resolving power, 3e6 AGC target, 50 ms maximum IT, 380 to 1,500 m/z, profile mode; up to 20 data-dependent MS2 scans per duty cycle, excluding charges 1 or 8 and higher, dynamic exclusion window 10 s, isolation window 0.7 m/z, fixed first mass 100 m/z, resolving power 60,000, AGC target 1e5 (min 1e3), maximum IT 100 ms, (N)CE 32.

Acquired raw files were searched in MaxQuant^51^ (1.6.17.0) against a *Drosophila melanogaster* fasta database downloaded from UniProtKB. Fixed cysteine modification was set to C6H11ON (+113.084064). Variable modifications were set to include Acetyl (protein N-term), Oxidation (M), Gln->pyroGlu, Deamidation (NQ) and Phospho (STY). Match-between-runs and second peptide search were set to active. All FDRs were set to 1 %. The output “evidence.txt” files were then re-processed in R using in-house scripts. Briefly, MS1 parent intensities were normalized per fraction; evidence reporter intensities were corrected using the relevant TMT lot’s purity table, then normalized using the Levenberg-Marquardt procedure and scaled to normalized parent MS1 intensity. Peptidoform intensity values were log_10_ transformed. The TMT/replicate-specific batch effect was corrected using Internal Reference Scaling, then values were re-normalized (Levenberg-Marquardt procedure). Protein groups were inferred from observed peptidoforms, and, for each group, its expression vector across samples was calculated by averaging the log_10_ intensity vectors across samples of individual unique and razor peptidoform, scaling the resulting relative profile vector to an absolute value reflecting the intensity level of the most intense peptidoform according to the best flyer hypothesis (phospho-peptides and their unmodified counterpart peptide were excluded). Peptidoform and protein group log_2_ ratios were calculated per replicate to the corresponding control (FlpND) sample. Statistical significance was tested with the limma package, performing both a moderated t-Test and an F-test (limma package) for all other genotypes against the FlpND genotype. The Benjamini-Hochberg procedure was applied to compute significance thresholds at various pre-agreed FDR level (up to 10, FDR thresholds calculated globally for the F-test). Regardless of the test, protein groups with a significant P-value were deemed to be regulated if their absolute log_2_ ratio was larger than the 90 % least extreme individual control to control log_2_ ratios.

### Immunohistochemistry and confocal imaging

For quantitative and qualitative analysis of ShakB localization, brains of 2-day-old male flies were dissected in cold PBS. The brains were fixed for 25 min in 4 % PFA/PBS, washed 2 h in PBS and then 4 times for 15 min in 0.3 % PBST, blocked in 10 % Donkey normal serum in 0.3 % PBST (Agrisera, AS10 1564) for 3 hours (all at RT), and incubated with primary antibodies diluted in 0.3 % PBST containing 5 % Donkey normal serum for 48 h at 4 °C. Samples were washed for 5 h in PBS at 4 °C and then 4 times for 15 min in 0.3 % PBST at RT, and incubated with secondary antibodies diluted in 0.3 % PBST containing 5 % Donkey normal serum for 12 h. Samples were washed again 4 times for 15 min in 0.3 % PBST and in PBS for 5 min at RT. The samples were mounted on glass microscope glasses with 0.12 mm-deep spacers in VECTASHIELD® mounting medium (Vector laboratories, H-1000).

For the analysis of neuronal morphology, the dissections and immunostainings were done as described above, with the difference that 2-to-5-day-old flies were used for experiments. The GFP signal was enhanced using anti-GFP primary antibodies in combination with AF594-conjugated secondary antibodies.

For trans-Tango-mediated transsynaptic tracing, the whole CNS of 10-to-15-day-old flies was dissected in cold PBS, fixed and washed as described above. To detect the postsynaptic partners of HS cells, CF594 anti-RFP antibodies were used. The samples were incubated with antibodies diluted in 0.3 % PBST containing 5 % Donkey normal serum overnight at 4 °C, washed and mounted as described above.

Antibodies and dilutions used in the IHC experiments: anti-shakB rabbit serum antibody (kind gift of Alexander Borst, Max Planck Institute for Biological Intelligence, Martinsried, Germany; 1:800), goat anti-GFP (Abcam, ab6673; 1:500), goat anti-RFP (Rockland, 200-101-379S; 1:500), rabbit anti-GFP (Thermo Fisher, A11122; 1:500), donkey anti-goat AF488 (Abcam, ab150129; 1:1000), donkey anti-goat AF594 (Thermo Fisher, A32758; 1:1000), donkey anti-rabbit AF594 (Thermo Fisher, A21207; 1:1000), CF594 rabbit anti-RFP (Biotium, 20422; 1:500).

Images of ShakB immunostainings and neurobiotin labeling were acquired on Zeiss LSM800 confocal microscope using 20X air objective (420650-9901-000) or 63X oil immersion objective (420782-9900-799). Images of HS cells for morphological analysis and of transsynaptic labeling were acquired on a Leica SP8 confocal microscope using 25X water immersion (15506374) or 20X air objective (11506517), respectively. The images were processed using Fiji software.

### Reconstruction of neuronal morphology

To label individual HS cells, we used 2-3 days-old male flies of the following genotype: *+; 20XUAS-SPARC2-I-mCD8::GFP/VT058487-p65.AD; VT000343-GAL4.DBD.* Only flies with single labeled HS cells were used. After IHC and imaging (see above), confocal z-stacks of individual HS cells were used for neuronal reconstruction using Neutube ^52^. The generated .swc files were loaded into Imaris software (Imaris9.3.1) as filaments using PylmarisSWC extension, implemented in Python. The diameter of each segment was manually readjusted based on the confocal images. Parameters of Filament dendrite length (sum) and Filament Bounding BoxAA for each neuron were normalized to the size of the optic lobe. For Sholl analysis, the step resolution was adjusted to the total length of the dendrite to obtain an equal amount of Sholl intersections for each cell type. Axes X and Y from Filament Bounding BoxAA were used to compute the dendritic field area.

### Transsynaptic tracing

To identify postsynaptic partners of HS cells, trans-Tango flies carrying a mutant and wild-type shakB alleles

*shakB*^2^*,UAS-myrGFP.QUAS-mtdTomato-3xHA/UAS-myrGFP.QUAS-mtdTomato-3xHA; trans-Tango*

and

*shakB^FlpStopD^,UAS-myrGFP.QUAS-mtdTomato-3xHA/UAS-myrGFP.QUAS-mtdTomato-3xHA; trans-Tango*

were crossed with HS-specific GAL4 driver line R81G07. This way, the genetic diversity and growth conditions between individuals were reduced due to the comparison of wild-type and mutant flies from the same progeny (siblings). The crosses were maintained at 18 °C. Male flies from the progeny were collected after eclosion and kept for another 14-16 days at 18 °C. Fly CNS were dissected and immuno-stained as described above. The variant of *shakB* allele of each dissected fly was identified using primers MiL-F and FRTspacer_3p_for for FlpD batch, and primers 5’-CACACCAACGCAACGGTTATATA-3’ and 5’-CGGCCCTGTGAATTGTGAAC-3’ with subsequent sanger sequencing for *shakB*^2^ batch.

### Analysis of isoform expression

Quantifying the expression of shakB isoforms was performed with Salmon^53^ The transcriptome of *Drosophila melanogaster* was indexed with decoys following the instructions of the Salmon documentation. Briefly, the files dmel-all-chromosome-r6.43.fasta.gz and dmel-all-transcript-r6.43.fasta.gz were downloaded from Flybase on December 16 2021. The chromosome sequences were used as decoy and the index was constructed with default k-mer length 31.

The RNA-seq data from^54^ was downloaded from ENA (European Nucleotide Archive) using accession numbers PRJNA658010. Each replicate was quantified independently using Salmon default parameters with options -lA to detect the library type automatically.

### Electrophysiology

1-day-old male flies were briefly anesthetized on ice and tethered using beeswax to a 3D-printed holder with a hole in the middle fitting the head and the thorax. The head of the tethered fly was passed through the hole and bent down to expose the head’s backside. The proboscis was fixed to the thorax with beeswax to avoid head movement. We cut out the cuticle on the backside of the head using a sterile needle (30 G). After the neuropil with LPTCs was exposed, we cut the transverse muscle and removed any fat excess. To digest the neurolemma and gain access to cell bodies, we used a brief 20 s treatment with protease solution (1 mg/ml Protease Type XIV in extracellular solution, Sigma Aldrich, P5147) at RT. The protease was washed off with an extracellular saline solution. The holder with the tethered fly was placed on the set-up under 40× water-immersion objective, and cell bodies of LPTCs were additionally cleaned under visual control using a low-resistance patch pipette (tip ∼4 μm) filled with extracellular saline solution.

Patch electrodes of 5-7 MΩ resistance (thin wall, filament, 1.5 mm, WPI, Florida, USA) were pulled on DMZ Zeitz-Puller (Zeitz-Instruments Vertriebs GmbH) and filled with intracellular solution. Using a Multiclamp 700B amplifier (Molecular Devices, Sunnyvale, USA), the signals were filtered at 4 kHz, digitized at 10 kHz, and recorded via a digital-to-analog converter (PCI-DAS6025, Measurement Computing, Massachusetts, USA) with Matlab (Vers.9.2.0.556344, MathWorks, Inc., Natick, MA). The recorded membrane potential was corrected for junction potential (12 mV).

The solutions used for *in vivo* whole-cell patch-clamp recording: extracellular saline solution (in mM): 103 NaCl, 3 KCl, 5 *N*-tris(hydroxymethyl) methyl-2-aminoethane-sulfonic acid, 10 trehalose, 10 glucose, 2 sucrose, 26 NaHCO3, 1 NaH2PO4, 1.5 CaCl2, and 4 MgCl2, adjusted to 275 mOsm, bubbled with 95 % O2/5 % CO2, and pH equilibrated around 7.3; the intracellular solution (in mM): 140 potassium aspartate, 10 HEPES, 1 KCl, 4 MgATP, 0.5 Na3GTP, and 1 EGTA, pH 7.26, adjusted to 265 mOsm. In most experiments, 0.5 % Neurobiotin was added to the intracellular solution.

### Neurobiotin cell filling and visualization

After the recordings were accomplished, cells were filled with Neurobiotin using a positive current of 1 nA for 10 min. After filling, the tissue was left in the recording bath for another 10 min for the dye to diffuse.

The heads of flies were fixed in 4 % PFA/PBS at RT for 1 hr, and washed 3 times for 15 min in PBS. The brains were dissected out of the head capsule in PBS. To visualize neurobiotin we used TSA-mediated streptavidin labeling, as described in ^55^, with some modifications. For that, fixed brains were incubated in 0.5 % PBST; 4 % NaCl solution containing 0.5 % avidin-biotinylated HRP complex (ABC) solution overnight at 4 °C. After the incubation, samples were washed 3 times for 15 min in PBS and incubated in 0.0001% biotin-tyramide (Sigma Aldrich, SML2135-50MG) and 0.003 % H_2_O_2_ in 0.05 M borate buffer, pH 8.5, for 2 h at RT. The samples were washed for 15 minutes in PBS and 3 times for 15 min in 0.3 % PBST. After washing, the samples were incubated Streptavidin-Alexa 546 (Thermo Fisher, S32356) or Streptavidin-Alexa 488 (Thermo Fisher, S32354) diluted 1:500 in 0.5 % PBST; 4 % NaCl solution at 4 °C overnight. The mounting and imaging of brains were performed as described above.

### Visual stimuli for electrophysiology

Visual stimuli were presented on the screen with the shape of a quarter sphere (diameter 4.6 cm) using an LED projector (Texas Instruments DLP LightCrafter Evaluation Module) at a frame rate of 60 Hz. A reflective filter (ND10A, Thorlabs) was positioned in front of the projector to block and reflect red light, which was then captured by a photodiode and used for synchronizing the stimulus and the recorded voltage responses of a cell. The stimulus presented on the screen had both green (peak of LED at ∼520 nm) and blue light (peak of LED at ∼470 nm). The scripts to present visual stimuli were written in Matlab using Psychtoolbox library^56^. To compensate for the spherical distortions of the screen, we created a customized lookup table that pre-deforms each frame on the GPU. The span of the visual field covered by the stimuli is ca. 140 ° horizontally and 85 ° vertically.

### Analysis of patch-clamp recordings

Cell voltage activity was recorded using the custom-designed software in LabView and analyzed in Matlab.

#### Full-field flashes

Full-field flashes were interleaved with intervals of dark screen in between. The average response per animal was computed across repetitions. The population average is the average of the responses of all flies with the same genetic background.

#### Gratings (contrast, direction and velocity selectivity)

To quantify the tuning of the cells to contrast, direction, and velocity we presented a moving grating stimulus with spatial frequency of 0.04 cycle/°, while changing one of the aforementioned parameters. The moving gratings were interleaved with intervals of the stationary gratings. The baseline computed during the stationary gratings was subtracted from the responses to the moving gratings. Population responses are averages of the responses of individual cells. For the direction tuning analysis, responses were normalized to the maximum response for each cell and then averaged across animals with the same genetic background.

#### Power spectrum analysis

For the power spectrum (PS) analysis, we first extracted the raw responses during the flash OFF periods and subtracted the global mean, such that the contribution of the lowest frequencies does not overshadow the contribution of higher frequencies. We used the Matlab fft function to compute the Fourier transform of the traces. The one-sided spectrum was normalized with respect to the bin size and squared to convert to the power of frequencies. Whenever multiple repetitions of the stimuli were available, we computed the PS for each repetition and averaged them to obtain the PS of the responses of one cell. The PS of each cell was normalized such that the total energy was 1. The PS of the population is the average PS of individual cells with the same genetic background. To quantify the contribution of three different ranges of frequencies, we subdivided the frequency range into two intervals: 0-10 Hz and 11-50 Hz. The sum of the power of the frequencies within these groups is depicted in Figure 4G as a percentage of the total power over all the frequencies to give the relative contribution of each interval.

#### Reconstruction of receptive field using a scanning rectangle stimulus

A 10 ° high and 2 ° wide rectangle scanned the visual field horizontally and vertically. Due to the long duration of the stimulus, a 60 s rolling window was used to subtract the mean voltage from the recording so that only the deviations from the baseline were used for the analysis. The responses were discretized into bins of 16.6 ms (duration of frame update). The position and direction of motion of the rectangle in each frame were weighted by the discretized voltage response to get a response vector at each point in the visual field. We computed the vector sum of the responses to the four cardinal directions and averaged this across multiple repetitions to compute the mean spatial receptive field of a cell which can be represented as a vector field. The location of each arrow in this vector field corresponds to a specific azimuth and elevation in the visual field of the fly; the direction represents the local preferred direction and the length indicates the relative strength of response. After normalizing the response vectors of each cell to the maximum, we averaged the responses of multiple cells from animals with the same genetic background to obtain the spatial receptive field of the population. In order to represent the deformations due to the spherical shape of the screen, we deform the receptive field to match the pre-deformed stimulus during the experiment. The horizontal/vertical profiles of the receptive field were computed using the length of the average response vector in the corresponding azimuth/elevation.

### Freely-walking arena

Individual flies walked freely on a 55 mm circular arena made of IR-transparent Perspex acrylic sheet with 3 mm tall walls. The walls were heated with an insulated nichrome wire to prevent the flies from walking on the walls and to encourage them to spend more time close to the center of the arena. The arena was covered with an IR-transparent acrylic sheet coated with Sigmacote^TM^ to prevent flies from walking on the roof. The entire behavioral setup is placed inside a custom-designed temperature-controlled compartment that maintains the internal temperature at 27 °C.

The visual stimulus was projected from the top on the outer face of the roof, which is covered with a projection screen (Gerriets OPERA^®^ Grey Blue Front and Rear Projection Screen). The stimulus was presented by an LED projector (Texas Instruments DLP LightCrafter Evaluation Module) at a framerate of 60Hz and pixel size of 12 px/mm. We used only the green LED (peak at ∼520 nm) of the projector since the relative sensitivity of the optomotor response of *Drosophila* has been shown to be highest for wavelengths between 350 and 500 nm ^57^

The fly was video recorded from the bottom using a monochrome USB3.0 camera (Flea3 FL3-U3-13YM) at a frame rate of 60 Hz and resolution 1024×1024 pixels. The arena was illuminated from the top by a custom-made panel of IR LEDs (850 nm). Since the projection screen, the roof and the arena are all made of IR-transparent material; the backlit flies appear as dark silhouettes on a bright background.

The stimuli (Figure 1A) were generated using a custom Python script and the various textures were made using the Psychopy library^58^. The stimulus was updated every frame depending on the position and orientation of the fly. We detected the contour of the fly after performing a pixel intensity thresholding. An ellipse was then fitted to this contour and position of the centroid and angle of the major axis of the fitted ellipse were used to determine the position and orientation of the fly. The delay between the fly movement and the update of the stimulus was 3 frame updates (<50 ms). The entire behavioral setup is placed inside a custom-designed temperature-controlled compartment that maintains the internal temperature at 27 ℃.

We presented a radial sinusoidal grating pattern, akin to a sinusoidal pinwheel, that was centered over the fly body. Rotating this pinwheel in either a clockwise or counterclockwise direction evoked a turning response, the optomotor response of the fly. The diameter of the pinwheel was kept 45 mm, since the optomotor response of the fly, which increased with an increase in the size of the pinwheel, saturated at 45 mm and did not change for larger pinwheels. An experiment session consisted of approximately 200 trials of 5 s pinwheel rotation preceded by 5 s without rotation. The direction, contrast and speed of rotation of the sinusoidal pinwheel were changed across trials. The contrast of the grating pattern was computed as:

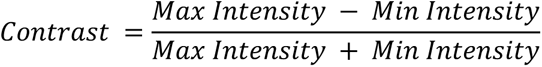

For experiments with unilateral motion, the pinwheel was divided into 2 segments and the direction of rotation of each segment (counterclockwise, clockwise or no rotation) was changed independently across trials to produce different combinations of rotational optic flow.

### Pre-processing of behavioral data

The position and orientation of the fly were determined for each frame during the experiment. The contour of the fly was extracted after performing a pixel intensity thresholding and an ellipse was fitted to this contour. The position of the centroid and the angle of the major axis of this fitted ellipse were used to determine the position and orientation of the fly, respectively. While this method is quick (a requirement for performing close-loop experiments) and works well for determining the orientation of the fly body, it does not provide the heading direction of the fly. The direction was estimated by processing the videos post-experiment. Otsu’s binarization was used to obtain two thresholds from the image, one for the body of the fly and one for the translucent wings. The direction of the line joining the center of mass (COM) of the body and the COM of the wings was used to determine the head direction of the fly, which was then used as the correct orientation for further analysis. All machine vision computations were performed using functions from the OpenCV Python library.

The speed and angular velocity of the fly were calculated from the change in position and orientation, respectively. In order to remove noise, we used a rolling median followed by a rolling average with a window length of 3 frames (∼50 ms). Events where the fly jumped were detected using a threshold of 100 mm/s or 1000 degrees/s. Trials in which the fly either jumped, went close to the walls of the arena (within 5 mm of the wall) or was inactive throughout the whole trial were excluded from further analysis.

### Estimation of saccades and path straightness

Fly locomotion is composed of long bouts of relatively straight motion interspersed with sharp turns, called saccades. We detected these saccades using a wavelet transform strategy inspired by Cruz et al. 2021^59^. The stationary wavelet transform (swt) of the angular velocity was computed using a biorthogonal 2.6 wavelet. The swt signal in the 10-20 Hz band was isolated to reconstruct the angular velocity data with an inverse stationary wavelet transformation. Peaks in this reconstructed angular velocity data that had a maximum angular speed higher than 200 degrees/s and with a width of more than 50 ms and less than 250 ms were deemed to be saccades.

In order to calculate the local curvature of locomotion, we first defined forward walking bouts as periods between two saccades that are longer than 333 ms (20 video frames) where the average speed of the fly is more than 5 mm/s. Then, a window of 333 ms was selected and centered around each point of a walking bout. The length of the straight line connecting the two endpoints of this window (the shortest distance between two points) and the perpendicular distance between this line and the midpoint of the actual trajectory (deviation from the shortest distance) were calculated. A straight walking bout will have a minimal deviation from a straight-line trajectory. As such, the straightness of a walking bout was defined as the ratio of the sum of the shortest distances and the sum of deviations from the shortest distance.

### Statistics

Details about statistical tests are provided in the figure legends and Supplementary Table 1.

### Material availability

Transgenic flies generated in this study are available on request.

### Data and software availability

Data used in the analysis will be uploaded to ISTA data repository and code used to generate the results is available on GitHub.

## Notes

### Competing Interest Statement

The authors have declared no competing interest.

