## Supplementary Material for "Gap junctions arbitrate binocular course control in flies"

### Supplementary information

### Supplementary Table 1. Detailed statistical analysis

| Figure | Sample Size | Statistical test | Statistical significance and effect size |
|---|---|---|---|
| 1J [prediction error] | CantonS - 13 | 1 sample T-test | total error P=1.651233e-06, t= 8.6618874<br>smooth error P=2.086168e-08, t=12.935533<br>saccadic error P=0.0012848829, t= 4.1763269 |
| 1L [smooth response] | CantonS - 17 | Mann-Whitney-Wilcoxon 2-sided | Full rotation vs. FtB P=1.0556E-06, U=272<br>BtF vs. FtB P=0.0005142, U=207<br>Full rotation vs. BtF P=0.000013983, U=217 |
| 1L [syn-saccadic response] | CantonS - 17 | Mann-Whitney-Wilcoxon 2-sided | Full rotation vs. FtB P=0.22753, U=170<br>BtF vs. FtB P=0.158792, U=155<br>Full rotation vs. BtF P=0.983416, U=113 |
| 1L [anti-saccadic response] | CantonS - 17 | Mann-Whitney-Wilcoxon 2-sided | Full rotation vs. FtB P=3.08861E-06, U=266<br>BtF vs. FtB P=3.75728E-06, U=236<br>Full rotation vs. BtF P=0.176681, U=79 |
| 2G [prediction error] | HS,VS>Kir2.1 - 21, UAS-Kir2.1 - 15 | Mann-Whitney-Wilcoxon 2-sided | VT058487-GAL4>UAS-Kir2.1 vs. UAS-Kir2.1 control;<br>total error P=0.000001486, U=308<br>smooth error P=0.000002395, U=305<br>saccadic error P=0.00009053, U=280 |
| 2J [smooth response] | HS,VS>Kir2.1 - 21, UAS-Kir2.1 - 15 | Mann-Whitney-Wilcoxon 2-sided | VT058487-GAL4>UAS-Kir2.1 vs. UAS-Kir2.1 control;<br>Full rotation P=0.002301, U=253<br>FtB P=0.01916, U=84<br>BtF P=0.2227, U=196 |
| 2J [syn-saccadic response] | HS,VS>Kir2.1 - 21, UAS-Kir2.1 - 15 | Mann-Whitney-Wilcoxon 2-sided | VT058487-GAL4>UAS-Kir2.1 vs. UAS-Kir2.1 control;<br>Full rotation P=0.004293, U=247<br>FtB P=0.01348, U=80<br>BtF P=0.07235, U=214 |
| 2J [anti-saccadic response] | HS,VS>Kir2.1 - 21, UAS-Kir2.1 - 15 | Mann-Whitney-Wilcoxon 2-sided | VT058487-GAL4>UAS-Kir2.1 vs. UAS-Kir2.1 control;<br>Full rotation P=0.158, U=113<br>FtB P=0.00000819, U=18<br>BtF P=0.0161, U=82 |
| 3F [immuno] | FlpND-6, FlpD-6, ShakB2-7, VT058487-6, FlpND_DB331-6, FlpD_DB331-6 | Mann-Whitney-Wilcoxon 2-sided | FlpND vs. FlpD P=0, U=144<br>FlpND vs. shakB2 P= 0.000017, U=168<br>FlpND vs. FlpNDxVT058487 P=0.001883, U=117<br>FlpND vs. FlpNDxDB331 P=0.000037, U=144 |
| 4D [resting Vm] | FlpND-15, FlpD-17, ShakB2-6, FlpNDxDB331-11 | Mann-Whitney-Wilcoxon 2-sided | |
| 4E [resting Vm variance] | FlpND-15, FlpD-17, ShakB2-6, FlpNDxDB331-11 | Mann-Whitney-Wilcoxon 2-sided | FlpND vs. FlpD P=1.620e-06, U=0 |
| 4F [power spectrum trace] | FlpND-10, FlpD-8, FlpNDxDB331-6, ShakB2-5 | | |

| | | | |
|---|---|---|---|
| 4G [power spectrum] | FlpND-10, FlpD-8, FlpNDxDB331-6, ShakB2-5 | Mann-Whitney-Wilcoxon 2-sided | FlpND vs. FlpD <10Hz P=0.002057, U=73<br>FlpND vs. FlpD 10-50Hz P=9.141e-05, U=1<br>FlpND-shakB2 <10Hz P=0.000666, U=50<br>FlpND-shakB2 10-50Hz P=0.000666, U=0 |
| 4H [ON and OFF response trace] | FlpND-10, FlpD-12, FlpNDxDB331-12 | | |
| 4I [ON edge response trace] | FlpND-8, FlpD-7, FlpNDxDB331-5 | | |
| 4J [OFF edge response trace] | FlpND-8, FlpD-7, FlpNDxDB331-5 | | |
| 4K [Direction Selectivity] | FlpND-10, FlpD-12, FlpNDxDB331-12 | Mann-Whitney-Wilcoxon 2-sided | FlpND vs. FlpD 180 deg P=0.0009, U=111<br>FlpND vs. FlpD 0 deg P=0.0059, U=95<br>FlpND vs. FlpD 45 deg P=0.0134, U=98<br>FlpND vs. FlpD 135 deg P=0.0003, U=115 |
| 4L [Contrast Tuning] | FlpND-12, FlpD-9, FlpNDxDB331-9 | Mann-Whitney-Wilcoxon 2-sided | FlpND vs. FlpD ND 25% P=0.0302, U=85<br>FlpND vs. FlpD ND 50% P=0.005, U=94<br>FlpND vs. FlpD ND 75% P=0.0012, U=100<br>FlpND vs. FlpD ND 100% P=0.0016, U=99 |
| 4M [Velocity Tuning] | FlpND-12, FlpD-12, FlpNDxDB331-8 | Mann-Whitney-Wilcoxon 2-sided | FlpND vs. FlpD PD 0.2 Hz P=0.0337, U=101<br>FlpND vs. FlpD ND 0.5 Hz P=0.0194, U=113<br>FlpND vs. FlpD ND 1 Hz P=0.0102, U=117<br>FlpND vs. FlpD ND 5 Hz P=0.0392, U=100<br>FlpND vs. FlpD ND 15 Hz P=0.0387, U=21 |
| 5A [FlpND HSN RFOF] | 5 | | |
| 5B [FlpND HSE RFOF] | 8 | | |
| 5C [FlpD HSN RFOF] | 5 | | |
| 5D [FlpD HSE RFOF] | 6 | | |
| 5G [size of receptive field] | FlpND-4, FlpD-4 | Mann-Whitney-Wilcoxon 1-sided | FlpND vs. FlpD HSN;<br>30% cutoff P=1.587e-02, U=2.300e+01<br>60% cutoff P=2.778e-02, U=2.200e+01 |
| 5H [HSN contralateral response/ipsilateral response] | FlpND-4, FlpD-4 | Mann-Whitney-Wilcoxon 2-sided | FlpND vs. FlpD HSN P=1.508e-01, U=2.000e+01 |
| 5I [size of receptive field] | FlpND-8, FlpD-6 | Mann-Whitney-Wilcoxon 1-sided | FlpND vs. FlpD HSE;<br>30% cutoff P=5.251e-01, U=2.400e+01<br>60% cutoff P=2.131e-02, U=4.000e+01 |
| 5J [HSE contralateral response/ipsilateral response] | FlpND-8, FlpD-6 | Mann-Whitney-Wilcoxon 2-sided | FlpND vs. FlpD HSE P=2.664e-03, U=4.600e+01 |

| | | | |
|---|---|---|---|
| 7G [prediction error] | FlpND-13, FlpD-11 | Mann-Whitney-Wilcoxon 2-sided | FlpND vs. FlpD ;<br>total error P=0.0005086, U=132<br>smooth error P=0.00377, U=122<br>saccadic error P=0.0007785, U=130 |
| 7J [smooth response] | FlpND-13, FlpD-11 | Mann-Whitney-Wilcoxon 2-sided | FlpND vs. FlpD;<br>Full rotation P=0.03506, U=380<br>FtB P=0.00000878, U=21<br>BtF P=0.3327, U=100 |
| 7J [syn-saccadic response] | FlpND-13, FlpD-11 | Mann-Whitney-Wilcoxon 2-sided | FlpND vs. FlpD;<br>Full rotation P=0.07521, U=44<br>FtB P=0.000201, U=44<br>BtF P=0.002229, U=45 |
| 7J [anti-saccadic response] | FlpND-13, FlpD-11 | Mann-Whitney-Wilcoxon 2-sided | FlpND vs. FlpD;<br>Full rotation P=0.7633, U=83<br>FtB P=0.002515, U=66<br>BtF P=0.01948, U=188 |
| S1A [path straightness] | CantonS - 17 | Mann-Whitney-Wilcoxon 2-sided | CantonS path straightness<br>Full rotation vs. no rotation P=1.747e-02, U=75<br>FtB vs. no rotation P=1.738e-04, U=35<br>BtF vs. no rotation P=1.761e-04, U=24 |
| S1B [saccade number] | CantonS - 17 | Mann-Whitney-Wilcoxon 2-sided | CantonS anti-saccade vs. syn-saccade<br>Full rotation P=9.889e-05, U=31<br>FtB P=7.040e-07, U=0<br>BtF P=2.696e-01, U=112 |
| S1D [saccade strength] | CantonS - 17 | Mann-Whitney-Wilcoxon 2-sided | CantonS anti-saccade vs. syn-saccade<br>P=1.667e-05, U=270 |
| S1D [saccade peak] | CantonS - 17 | Mann-Whitney-Wilcoxon 2-sided | CantonS anti-saccade vs. syn-saccade<br>P=3.564e-02, U=206 |
| S2A [path straightness] | HS,VS>Kir2.1 - 21, UAS-Kir2.1 - 15 | Mann-Whitney-Wilcoxon 2-sided | path straightness UAS-Kir2.1 vs. HS,VS-Kir2.1;<br>Full rotation P=1.503e-01, U=118<br>FtB P=4.711e-07, U=315<br>BtF P=4.673e-01, U=141<br>no rotation P=8.050e-02, U=108 |
| S2B [saccade number] | HS,VS>Kir2.1 - 21, UAS-Kir2.1 - 15 | Mann-Whitney-Wilcoxon 2-sided | UAS-Kir2.1 anti-saccade vs. syn-saccade;<br>Full rotation P=0.00004806, U=211<br>FtB P=0.000003392, U=0<br>BtF P=0.0745, U=156<br>HS,VS>Kir2.1 anti-saccade vs. syn-saccade;<br>Full rotation P=0.497, U=248<br>FtB P=0.002758, U=101<br>BtF P=0.1249, U=282 |
| S2D [saccade strength] | HS,VS>Kir2.1 - 21, UAS-Kir2.1 - 15 | Mann-Whitney-Wilcoxon 2-sided | UAS-Kir2.1 anti-saccade vs. syn-saccade;<br>P=1.605e-05, U=217<br>HS,VS>Kir2.1 anti-saccade vs. syn-saccade;<br>P=1.349e-05, U=394<br>syn-saccade UAS-Kir2.1 vs. HS,VS>Kir2.1<br>P=8.466e-04, U=262<br>anti-saccade UAS-Kir2.1 vs. HS,VS>Kir2.1 |

| | | | |
|---|---|---|---|
| | | | P=4.747e-03, U=246 |
| S2D [saccade peaks] | HS,VS>Kir2.1 - 21, UAS-Kir2.1 - 15 | Mann-Whitney-Wilcoxon 2-sided | UAS-Kir2.1 anti-saccade vs. syn-saccade;<br>P=1.620e-03, U=189<br>HS,VS>Kir2.1 anti-saccade vs. syn-saccade;<br>P=3.248e-03, U=338<br>syn-saccade UAS-Kir2.1 vs. HS,VS>Kir2.1<br>P=1.315e-01, U=205<br>anti-saccade UAS-Kir2.1 vs. HS,VS>Kir2.1<br>P=2.271e-02, U=229 |
| S5D [sholl HSN] | wild-type - 9<br>FlpD - 4<br>shakB2 - 5 | | |
| S5E [sholl HSE] | wild-type - 6<br>FlpD - 2<br>shakB2 - 4 | | |
| S5F [sholl HSS] | wild-type - 7<br>FlpD - 4<br>shakB2 - 4 | | |
| S5G [HSN dendrite length] | wild-type - 9<br>FlpD - 5<br>shakB2 - 5 | Mann-Whitney-Wilcoxon 2-sided | wild-type vs. FlpD<br>P=0.6993, U=19<br>wild-type vs. shakB2<br>P=0.5443, U=22 |
| S5H [HSN dendrite area] | wild-type - 10<br>FlpD - 4<br>shakB2 - 5 | Mann-Whitney-Wilcoxon 2-sided | wild-type vs. FlpD<br>P=0.2398, U=29<br>wild-type vs. shakB2<br>P=0.5941, U=30 |
| S5I [HSE dendrite length] | wild-type - 7<br>FlpD - 2<br>shakB2 - 4 | Mann-Whitney-Wilcoxon 2-sided | wild-type vs. FlpD<br>P=0.8889, U=10<br>wild-type vs. shakB2<br>P=0.5273, U=8 |
| S5J [HSE dendrite area] | wild-type - 6<br>FlpD - 2<br>shakB2 - 4 | Mann-Whitney-Wilcoxon 2-sided | wild-type vs. FlpD<br>P=0.0714, U=0<br>wild-type vs. shakB2<br>P=0.1143, U=4 |
| S5K [HSS dendrite length] | wild-type - 9<br>FlpD - 4<br>shakB2 - 4 | Mann-Whitney-Wilcoxon 2-sided | wild-type vs. FlpD<br>P=0.7105, U=21<br>wild-type vs. shakB2<br>P=0.6042, U=22 |
| S5L [HSS dendrite area] | wild-type - 8<br>FlpD - 4<br>shakB2 - 4 | Mann-Whitney-Wilcoxon 2-sided | wild-type vs. FlpD<br>P=0.3677, U=22<br>wild-type vs. shakB2<br>P=0.2411, U=16 |
| S7A [PD response] | FlpND - 10<br>CantonS - 6 | Mann-Whitney-Wilcoxon 2-sided | FlpND vs. CantonS P=5.622e-01, U=36 |
| S7B [PD response] | FlpND - 10<br>CantonS - 6 | Mann-Whitney-Wilcoxon 2-sided | FlpND vs. CantonS P=7.268e-02, U=13 |
| S7D [ON flash response] | FlpND- 10<br>FlpD-8<br>FlpNDxDB331-6 | Mann-Whitney-Wilcoxon 2-sided | FlpND vs. FlpD P=4.598e-01, U=4.900e+01<br>FlpND vs. FlpND,UAS-Flp,DB331-Gal4 P=7.268e-02, U=4.700e+01 |
| S7D [OFF flash response] | FlpND- 10<br>FlpD-8<br>FlpNDxDB331-6 | Mann-Whitney-Wilcoxon 2-sided | FlpND vs. FlpD P=6.216e-03, U=7.000e+01<br>FlpND vs. FlpND,UAS-Flp,DB331-Gal4 P=1.471e-01, U=4.400e+01 |

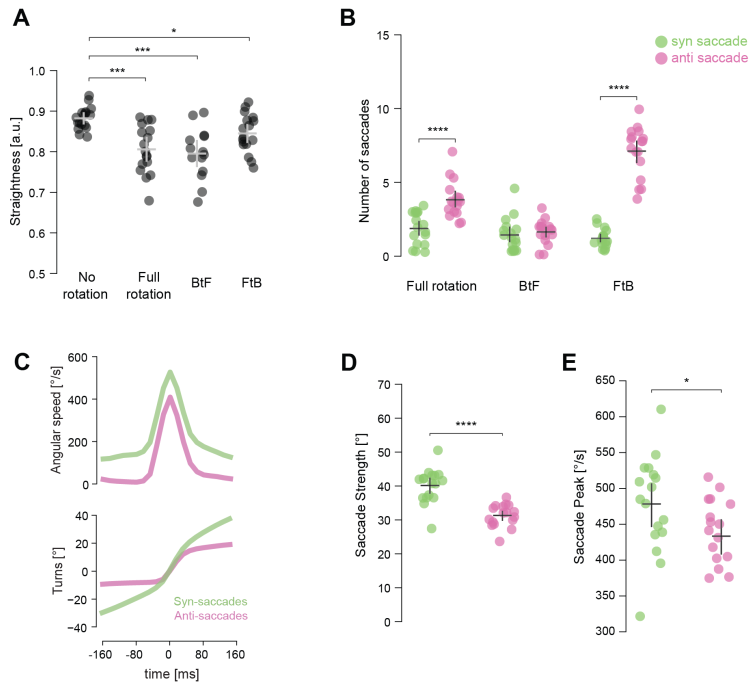


**Figure S1. Path straightness and properties of saccades for CantonS flies**

(A) Path straightness of CantonS flies in response to full-field, BtF and FtB rotation.

(B) Number of syn- and anti-saccades per trial for CantonS flies.

(C) Mean angular speed (left) and turns (right) during a syn-saccade and an anti-saccade.

(D) Total turns made per saccade (in a ~160 ms window centered on the saccade peak) for syn- and anti-saccades by CantonS flies.

(E) Maximum angular speed during a saccade for syn- and anti-saccades for CantonS flies.

Mann-Whitney U test was applied in all panels, $*p < 0.05$, $**p < 0.01$, $***p < 0.001$, $****p < 0.0001$. The number of flies and exact *p*-values for each experiment are listed in Suppl. Table1.

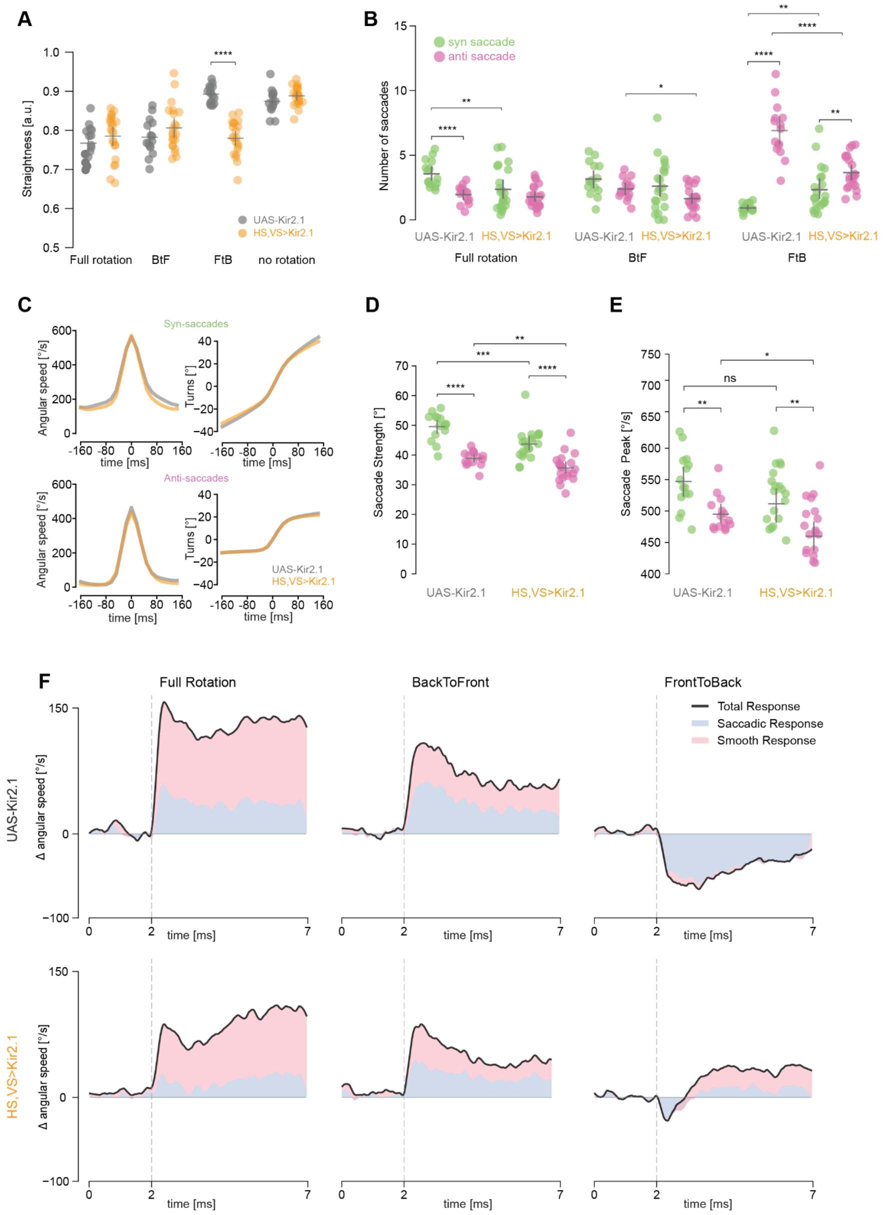


**Figure S2. Path straightness and properties of saccades for UAS-Kir2.1 control and HS & VS > Kir2.1 flies**

(A) Path straightness of UAS-Kir2.1 control and HS & VS silenced flies in response to full-field, BtF and FtB rotation.

(B) Number of syn- and anti-saccades per trial for UAS-Kir2.1 control and HS & VS silenced flies.

(C) Mean angular speed (left) and turns (right) during a syn-saccade and an anti-saccade.

(D) Total turns made per saccade (in a 160 ms window centered on the saccade peak) for syn- and anti-saccades in UAS-Kir2.1 control and HS & VS silenced flies.

(E) Maximum angular speed during a saccade for syn- and anti-saccades.

(F) Stacked plot showing mean angular speed and the contribution of smooth and saccadic turning.

Mann-Whitney U test was applied in all pannels, $*p < 0.05$, $**p < 0.01$, $***p < 0.001$, $****p < 0.0001$. The number of flies and exact *p*-values for each experiment are listed in Suppl. Table1.

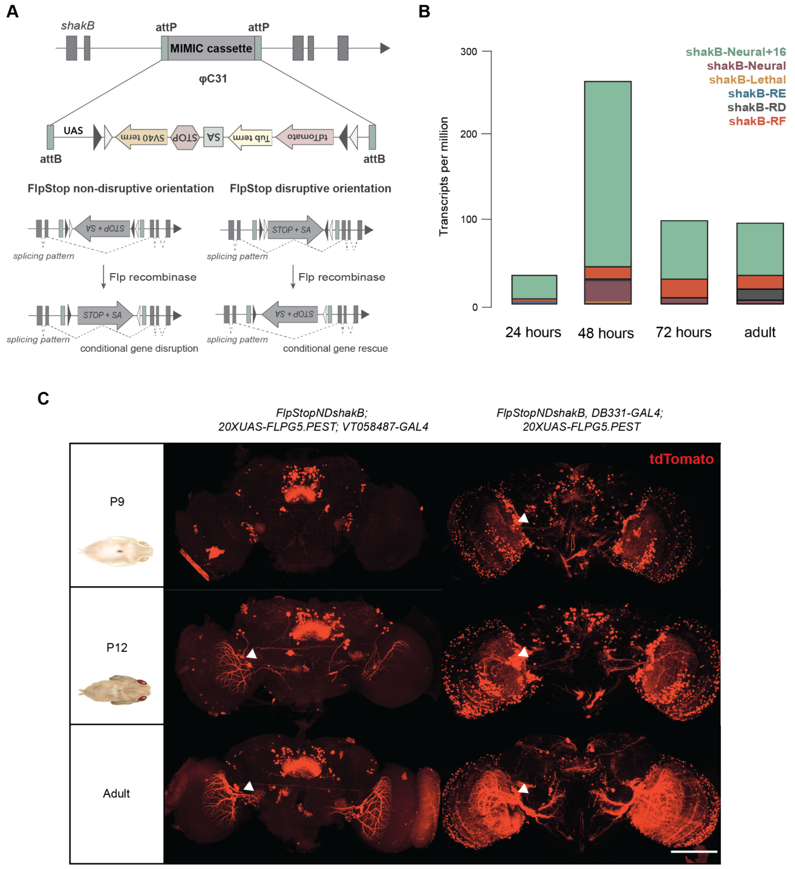


**Figure S3. FlpStop technique for disruption of the *shakB* gene**

(A) Schematic of the structure of the FlpStop cassette and of the gene disruption mechanisms[23]. The cassette is integrated into MiMIC insertion using PhiC31-integrase. The FlpStop cassette contains a splice acceptor (SA), transcriptional terminators (Tub α 1 terminator and the SV40), and stop codons in three frames (STOP). The cassette can be integrated into disrupting (FlpStopD) or non-disrupting (FlpStopND) orientation. The disrupting orientation can be used for a constitutive gene knock-out, while the non-disrupting orientation can trigger cell-specific gene inactivation in combination with flippase (Flp) and a GAL4-driver line.

(B) The expression of shakB isoforms in the optic lobe of *Drosophila* throughout pupal development. The analysis was performed on single-cell sequencing data of transcriptomes in the developing *Drosophila* visual system (see Methods). Isoform shakB-Neural+16 comprises shakB-RH, shakB-RI, shakB-RG; shakB-Neural corresponds ro shakB-RC, and shakB-Lethal to shakB-RA.

(C) The dynamics of the inversion of the FlpStop cassette using two distinct LPTC-specific driver lines - DB331-Gal4 and VT058487-Gal4. The expression of the tdTomato was used as a marker of the cassette inversion. The onset of the cassette inversion is around P9 for DB331-Gal4 and around P12 for VT058487-Gal4, somas of LPTCs are indicated with white arrows. Scale bar: 100 µm.

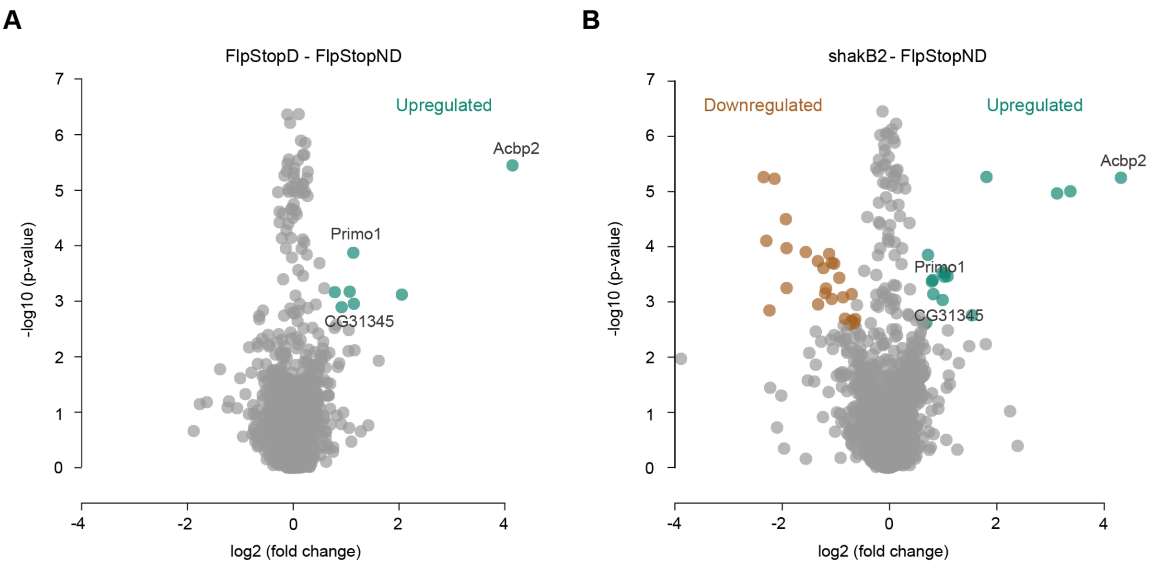


**Figure S4. Proteomic analysis of FlpD and *shakB*$^{2}$ mutant brains**

Total protein lysates obtained from fly brains were analyzed through liquid chromatography mass-spectrometry (LC-MS)

(A) Protein level quantification (fold change) and statistical significance assessment (p-value) for FlpD mutant model were performed against FlpND control flies. Proteins that showed significant changes in expression levels are depicted in brown (down-regulated) and green (up-regulated).

(B) The same for the *shakB2* mutant model.

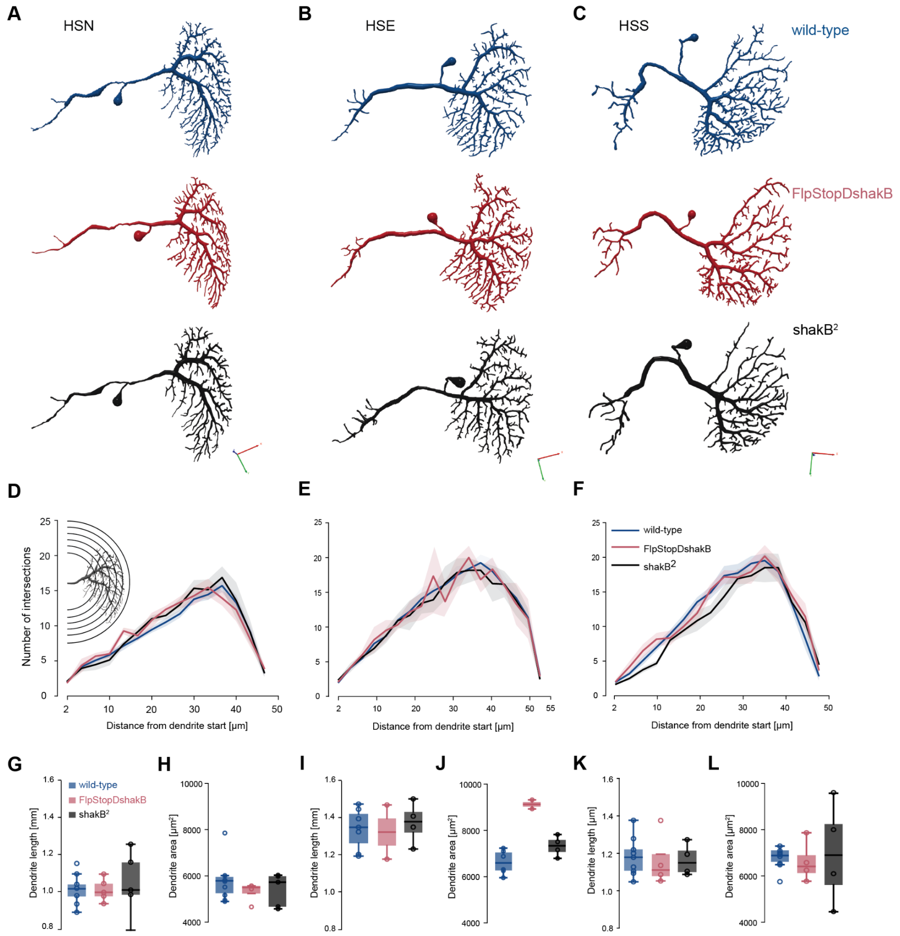


**Figure S5. Loss of gap junctions does not affect the morphology of dendrites in HS cells**
(A-C) Examples of reconstructed HSN, HSE and HSS cells for every genotype.
(D-F) Sholl intersection profile of HS dendrites in the wild type and two mutant lines (mean ± SEM). The number of intersections for each HS type was equalized between genotypes.
(G, I, K) Total length of HSN, HSE and HSS dendrites in the wild type and two mutant lines.
(H, J, L) Dendritic field area for HSN, HSE and HSS in the wild type and two mutant lines.

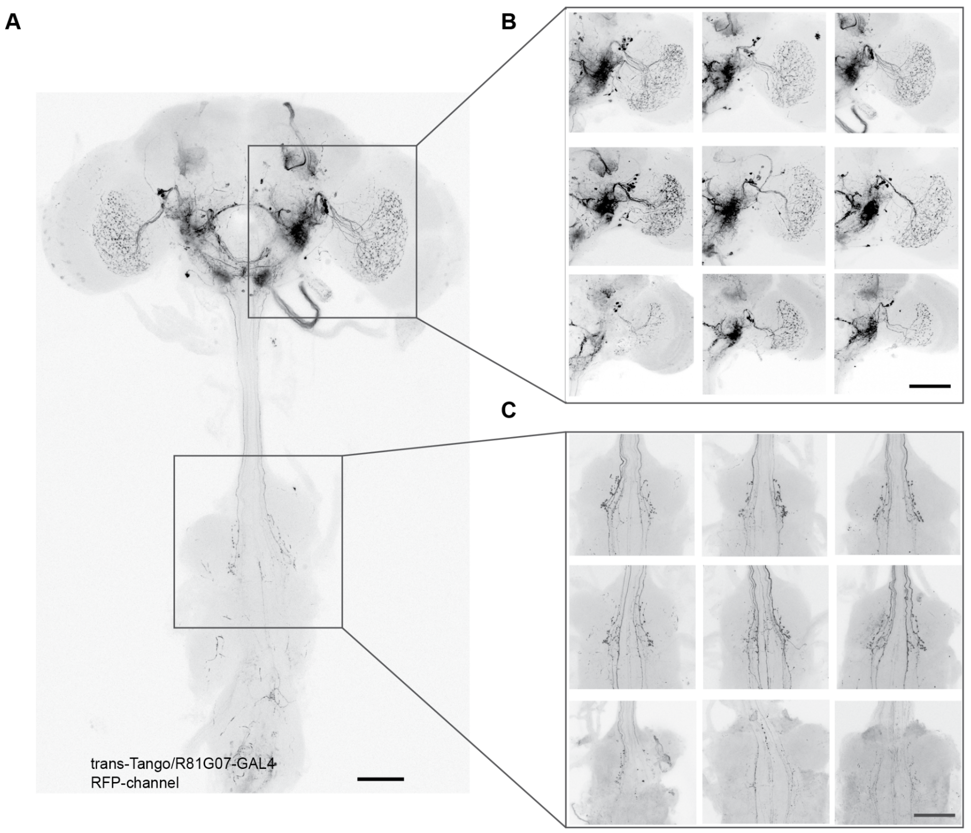


**Figure S6. Trans-synaptic labeling does not reveal the loss of chemical postsynaptic partners of HS cells in FlpD flies**

(A) Expression pattern of the postsynaptic marker (tdTomato) in HS>trans-Tango flies.

(B) Examples of labeled synaptic partners of HS cells in the optic lobe and the posterior slope in the wild type and two mutant lines.

(C) Examples of labeled synaptic partners of HS cells in the VNC in the wild type and two mutant lines. Scale bar: 50 µm.

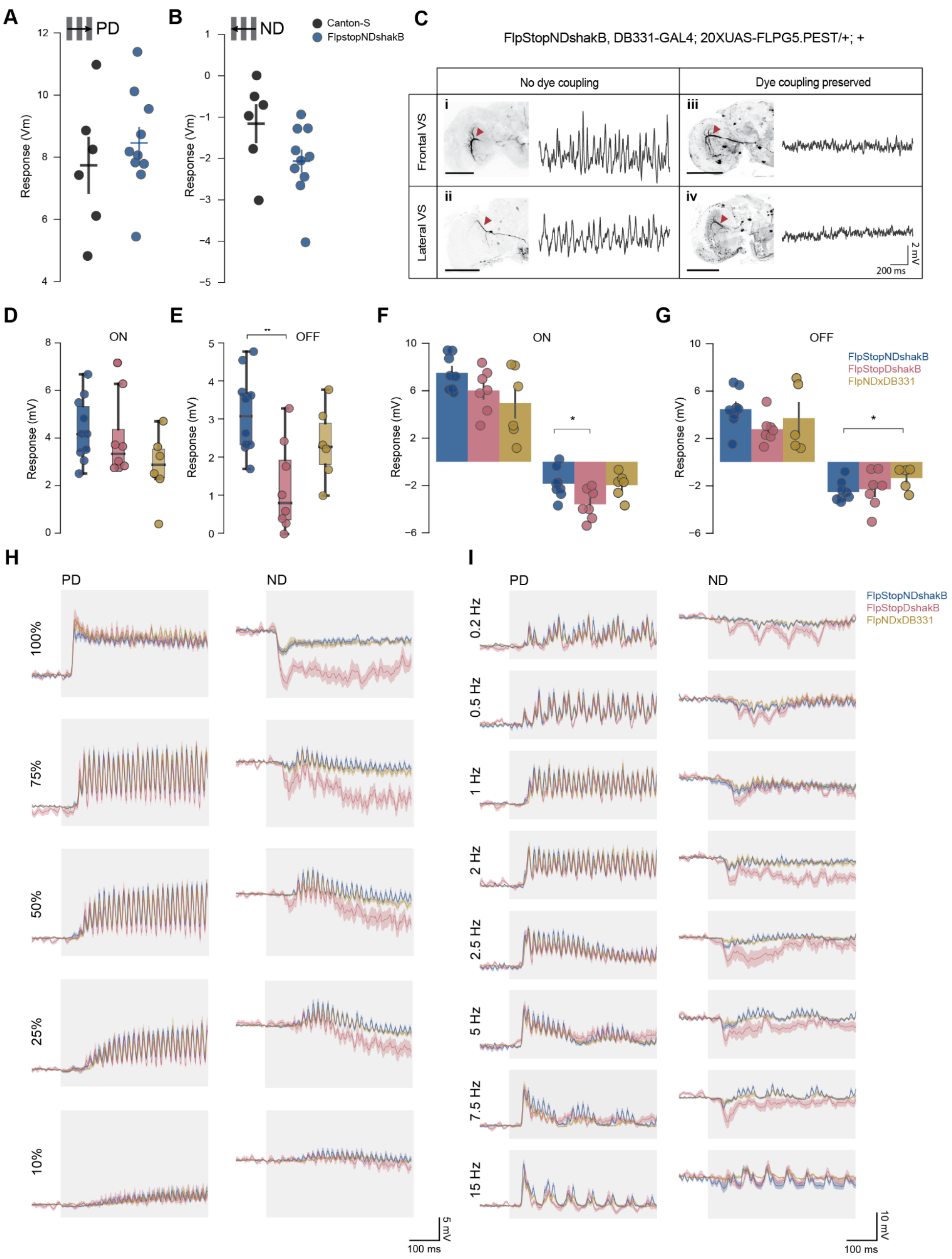


**Figure S7. Additional characterization of FlpStop-induced *shakB* inactivation on physiological properties of LPTCs**

(A) Average voltage changes of HS cells in Canton-S and FlpND flies during 2 s presentation of square-wave gratings moving in PD (mean ± SEM).

(B) Same as in (A) but for null direction (ND).

(C) Fast membrane oscillations of LPTCs are cell-intrinsic. Example traces of membrane potential and neurobiotin coupling of VS cells in flies with LPTC-specific inactivation of ShakB protein. In contrast to VS cells preserving neurobiotin coupling (iii, iv), VS cells lacking neurobiotin coupling with other LPTC cells (i, ii) exhibit fast membrane fluctuations. Scale bar: 50 µm, red triangle indicates injected/recorded cell.

(D) Voltage response of HS cells in wild type and mutant flies to full-field light ON flash during 50 ms onset of the stimulus. Oscillations are a stimulus artifact from the projector's refresh rate.

(E) Same as (D) but for light OFF flash stimulus.

(F) Average voltage responses of HS cells in wild-type and mutant flies to drifting ON-edges moving in PD and ND.

(G) Same as (F) but for OFF-edge.

(H) Average voltage response traces of HS cells in wild type and mutant flies to gratings with different contrast moving in PD and ND at a temporal frequency of 1 Hz (mean± SEM).

(I) Average voltage response traces of HS cells in wild type and mutant flies to gratings moving with different temporal frequency in PD and ND (mean± SEM).

For (A, B, D-G) Mann-Whitney U test was applied, $*p < 0.05$, $**p < 0.01$.

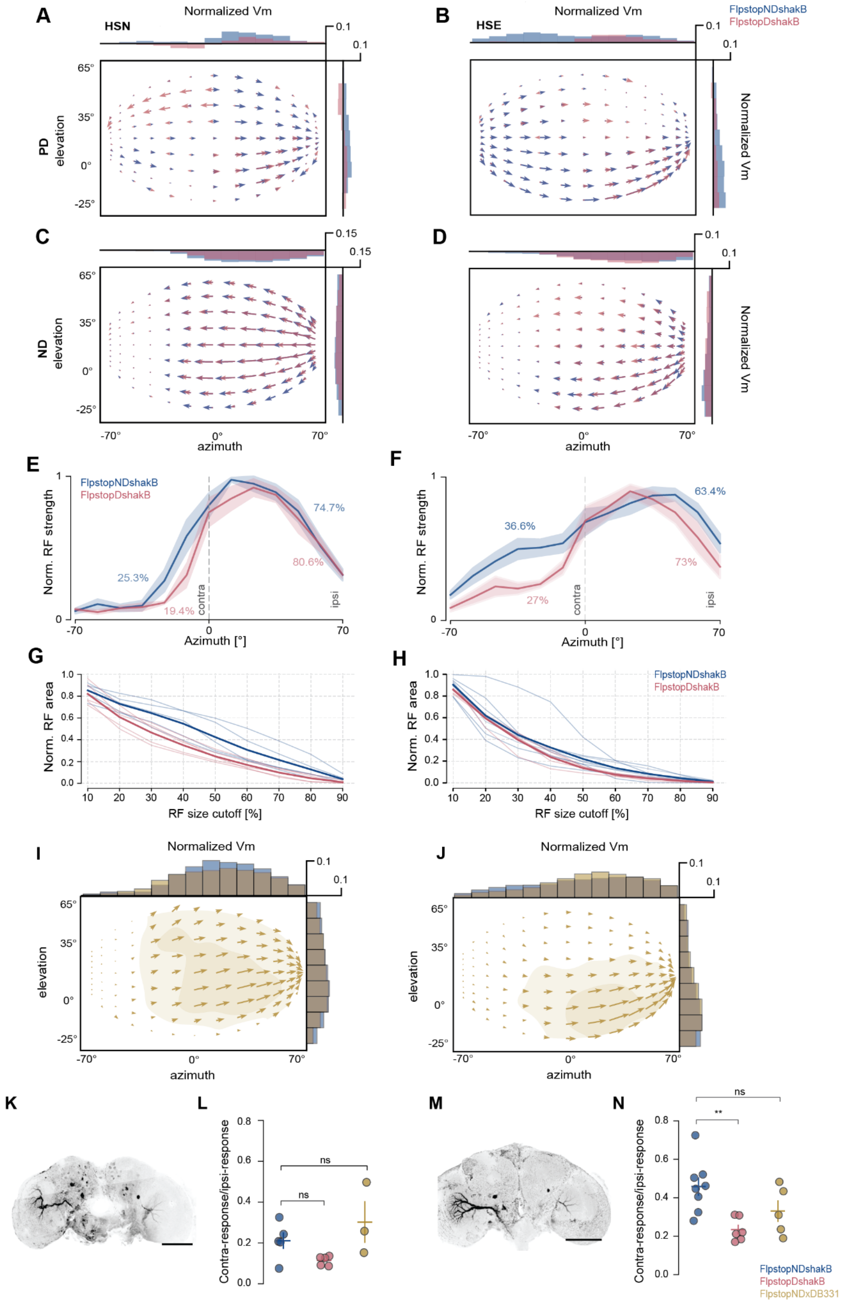


**Figure S8. Additional characterization of FlpStop-induced *shakB* inactivation on the receptive fields (RF) of HS cells**

(A-B) The pattern of local motion sensitivity of (A) HSN and (B) HSE cells for FlpND and FlpD flies to PD motion. Horizontal and vertical bar plots show average response along azimuth and elevation, respectively. (C, D) Same as (A, B) but for local motion in ND.

(E) Normalized strength of responses of HSN neurons across the azimuth, computed as the sum of all local response vectors at a given azimuth.

(F) Same as (E) but for HSE cells.

(G) Size of RF of HSN cells (as a proportion of total recorded visual field) for different cutoffs of the maximal response.

(H) Same as (G) but for HSE neurons.

(I) Spatial RF reconstructed from the responses to local motion stimulus of HSN cells in flies with induced inversion of FlpStop-cassette in LPTCs (*FlpND,DB331-GAL4; UAS-Flp/+;+*). Light-shaded and dark-shaded areas represent 30% and 60% of the maximal strength of the response. Horizontal and vertical bar plots show response along azimuth and elevation respectively for FlpND and induced mutant flies.

(J) Same as (I), for HSE cells.

(K) An example of neurobiotin injection into an individual HSN cell in *FlpND,DB331-GAL4; UAS-Flp* flies. Scale bar 100 µm.

(L) Relative strength between the contralateral and ipsilateral visual fields of individual HSN cells. Bars: mean ± SEM.

(M) Same as (K) but for an HSE neuron.

(N) Same as (L) but for HSE neurons.

Mann-Whitney U test was applied in (L) and (N), $*p < 0.05$, $**p < 0.01$. The number of cells and exact *p*-values for each experiment are listed in Suppl. Table1.

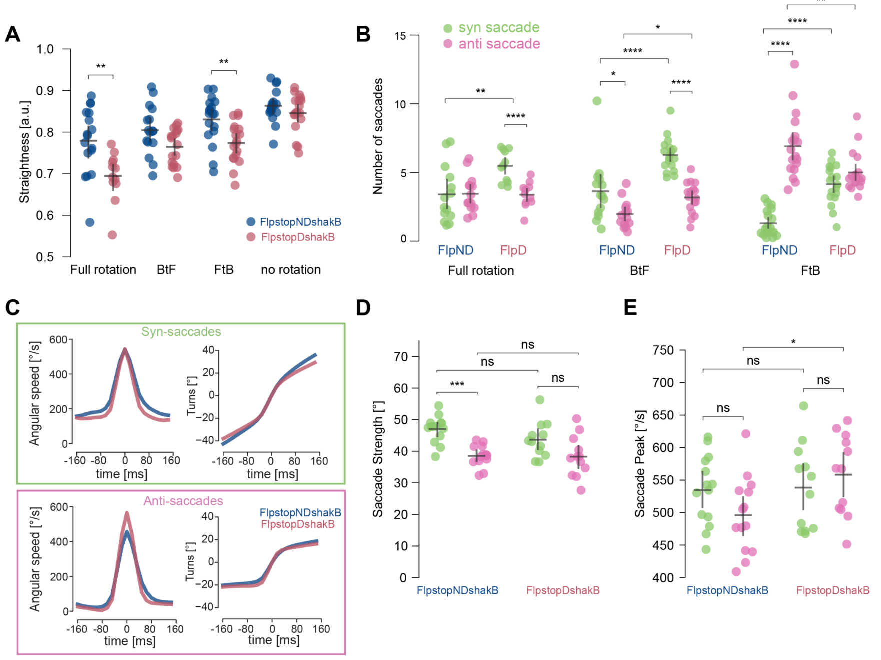


**Figure S9. Path straightness and properties of saccades for FlpND and FlpD flies**

(A) Path straightness for FlpND and FlpD flies in response to full-field, BtF and FtB rotation.
(B) Number of syn- and anti-saccades per trial for FlpND and FlpD flies.
(C) Mean angular speed (left) and turns (right) during a syn-saccade and an anti-saccade.
(D) Total turns made per saccade (in a 160 ms window centered on the saccade peak) for syn- and anti-saccades in FlpND and FlpD flies.
(E) Maximum angular speed during a saccade for syn- and anti-saccades in FlpND and FlpD flies.
Mann-Whitney U test was applied in all panels, $*p < 0.05$, $**p < 0.01$, $***p < 0.001$, $****p < 0.0001$. The number of flies and exact *p*-values for each experiment are listed in Suppl. Table1.
